# Structural basis of TRPC4 regulation by calmodulin and pharmacological agents

**DOI:** 10.1101/2020.06.30.180778

**Authors:** Deivanayagabarathy Vinayagam, Dennis Quentin, Oleg Sitsel, Felipe Merino, Markus Stabrin, Oliver Hofnagel, Maolin Yu, Mark W. Ledeboer, Goran Malojcic, Stefan Raunser

## Abstract

Canonical transient receptor potential channels (TRPC) are involved in receptor-operated and/or store-operated Ca^2+^ signaling. Inhibition of TRPCs by small molecules was shown to be promising in treating renal diseases. In cells, the channels are regulated by calmodulin. Molecular details of both calmodulin and drug binding have remained elusive so far. Here we report structures of TRPC4 in complex with a pyridazinone-based inhibitor and a pyridazinone-based activator and calmodulin. The structures reveal that both activator and inhibitor bind to the same cavity of the voltage-sensing-like domain and allow us to describe how structural changes from the ligand binding site can be transmitted to the central ion-conducting pore of TRPC4. Calmodulin binds to the rib helix of TRPC4, which results in the ordering of a previously disordered region, fixing the channel in its closed conformation. This represents a novel calmodulin-induced regulatory mechanism of canonical TRP channels.

## INTRODUCTION

Transient receptor potential (TRP) ion channels mediate a plethora of vital cellular functions, including nociception, mechanosensation and store-operated Ca^2+^ signaling (Clapham, 2003). Members belonging to the canonical TRP subfamily (TRPC) are involved in neuronal development and plasticity, as well as in vasorelaxation and kidney dysfunction (Hall et al., 2019; Kochukov et al., 2012; Phelan et al., 2013; Riccio et al., 2009). As a result, malfunction is often linked to pathologies such as neurological disorders and cardiac hypertrophy (Selvaraj et al., 2010; Wu et al., 2010). This class of non-selective cation channels can be further subdivided into TRPC1/4/5, TRPC2 (which is a pseudogene in humans) and TRPC3/6/7 groups, based on their sequence similarity. Within the TRPC1/4/5 sub-group, TRPC4 and TRPC5 share the highest sequence identity of 70% (Plant and Schaefer, 2003). Both proteins can form homo-tetrameric channels that allow the passage of Ca^2+^, but also Na^+^ ions to a lesser extent (Minard et al., 2018; Owsianik et al., 2006). In contrast, TRPC1, which shares approximately 48% identity with TRPC4/5, rather participates in the formation of hetero-tetrameric TRPC1/4/5 channels (Bröker Lai et al., 2017). Whether or not functional homo-tetrameric TRPC1 channels exist *in vivo* and what their potential physiological impact may be, is currently not known.

TRPC4 is widely expressed in various tissues associated with the nervous-, cardiovascular- and immune system (Freichel et al., 2014). It has been shown to be necessary for neurite outgrowth and its expression is upregulated in axonal regeneration after nerve injury (Wu et al., 2007). Channel activation results in a depolarization of the cell membrane, followed by a surge of intracellular Ca^2+^ levels. The regulation of the activity of TRPC channels, however, is multi-faceted and ranges from modulation by endogenous and dietary lipids to surface receptors, the redox environment and various types of cations (Jeon et al., 2012). Even within the TRPC1/4/5 subgroup, regulatory mechanisms can differ and are dependent on the respective cellular environment in combination with the experimental method used for the measurement (Plant and Schaefer, 2003). TRPC4 interacts with multiple proteins that can modulate its activity. This includes the ER-resident calcium sensor Stim1, the lipid binding protein SESTD1 and G-protein G_αi2_ (Jeon et al., 2012; Lee et al., 2010; Miehe et al., 2010; Zeng et al., 2008).

Several studies also established a role for TRPC4 as a store-operated channel (SOC) (Wang et al., 2004; Warnat et al., 1999). Here, two key proteins, calmodulin (CaM) and inositol 1,4,5-triphosphate receptor (IP_3_R), compete for the same binding site on TRPC4 (Mery et al., 2001; Tang et al., 2001). First, Ca^2+^-dependent binding of CaM inhibits the channel in the resting state. When intracellular Ca^2+^ levels decrease, the activating IP_3_ receptor directly interacts with TRPC4, displacing CaM, to restore channel activity (Kanki et al., 2001). This process, which is also known as conformational coupling, represents a primary regulation mechanism of gating for SOCs (Berridge and Berridge, 2004). However, a detailed mechanistic understanding of CaM inhibition or IP_3_R activation remains elusive.

Due to their implication in various diseases, TRPC channels also constitute a prime target for pharmacological intervention by small molecules (Minard et al., 2018). Activation of channels by the natural compound (-)-Englerin A (EA), which shows high potency and selectivity for TRPC4/5, inhibits tumor growth of renal cancer cells through increased Ca^2+^ influx (Akbulut et al., 2015; Carson et al., 2015). Other activators include riluzole, BTD and the glucocorticoid methylprednisolone (Beckmann et al., 2017; Richter et al., 2013). However, these compounds are typically either less potent or show varying specificity.

Inhibitors of TRPC4/5 are mostly used to target renal diseases such as focal segmental glomerulosclerosis (FSGS) (Mundel et al., 2019; Zhou et al., 2017), but can also have a therapeutic effect on the central nervous system (CNS) (Just et al., 2018; Yang et al., 2015). Currently, two compounds are in clinical trials aiming to treat a proteinuric kidney disease and anxiety disorder/depression, conducted by Goldfinch Bio (NCT03970122) and Hydra/Boehringer Ingelheim (NCT03210272), respectively (Mundel et al., 2019; Wulff et al., 2019). While xanthine-based inhibitors, such as HC-070 and HC-608 (formerly known as Pico145) have assisted in advancing the field of pharmacological modulation of TRPC1/4/5 due to their exceptional high potency, they suffer from poor physiochemical properties such as low solubility (Just et al., 2018; Rubaiy et al., 2017).

Recently, a novel class of small molecule modulators selective for TRPC4/5 was identified in a high-throughput screen, building up on a piperazinone/pyridazinone scaffold (Yu et al., 2019). Among this class of modulators are the activator GFB-9289 and the inhibitor GFB-8438. In particular, GFB-8438 showed promise as a potential drug for the treatment of proteinuric kidney disease, exhibiting overall favorable *in vitro* and *in vivo* properties (Yu et al., 2019). *In vitro*, mouse podocytes were protected from protamine-induced injury when treated with the inhibitor. Importantly, GFB-8438 also demonstrated robust efficacy in a hypertensive deoxycorticosterone acetate (DOCA)-salt rat model of FSGS, in which both albumin concentration and total protein levels were significantly reduced (Yu et al., 2019). However, information on the TRPC4/5 binding site and the mode-of-action of this novel compound class are still lacking. To date, the structures of TRPC6 in complex with the activator AM-0883 and the inhibitor AM-1473 are the only source of information regarding how modulation of TRPC channels is mediated on a molecular scale (Bai et al., 2020). Although insights gained from different apo structures of TRPC4/5 have advanced our understanding of this medically important TRPC subfamily (Duan et al., 2018; Vinayagam et al., 2018), a molecular understanding of pharmacological modulation by small molecules as well as key regulatory proteins such as CaM and IP_3_R remains unknown. Here we report four cryo-EM structures of TRPC4 in its apo form and in complex with the inhibitor GFB-8438, the activator GFB-9289 and CaM, respectively. Based on the analysis of the structures we propose mechanistic pathways by which CaM and small molecules exert their action to modulate the activity of the channel.

## Results and discussion

### Cryo-EM structures of full-length TRPC4 in complex with a small molecule inhibitor and activator

We previously reported the high-resolution apo structure of zebrafish TRPC4 in amphipols in its closed state (Vinayagam et al., 2018). To understand how channel activity is modulated by pharmacological compounds, we examined the complex of TRPC4 with the inhibitor GFB-8438 and the activator GFB-9289. Both compounds belong to the same novel class of TRPC4/5-selective modulators, which contain a common piperazinone/pyridazinone core. We first conducted fluorescent dye based Ca^2+^ uptake assays on TRPC4-transfected HEK293T cells to confirm their inhibitory or activating effects. GFB-8438 indeed acted as inhibitor of TRPC4, whereas GFB-9289 specifically activated Ca^2+^ influx by TRPC4 (Figure S1A).

We then formed the TRPC4 complexes with the respective small molecules. We did not add cholesteryl hemisuccinate and exogenous lipid molecules during purification to exclude potential interference with ligand binding. Using cryogenic electron microscopy (cryo-EM) and single particle analysis, we then determined the structures of GFB-8438-bound and GFB-9289-bound TRPC4 to an average resolution of 3.6 Å and 3.2 Å, respectively (Figure 1, 2, Figure S2, S3).

**Figure 1.**
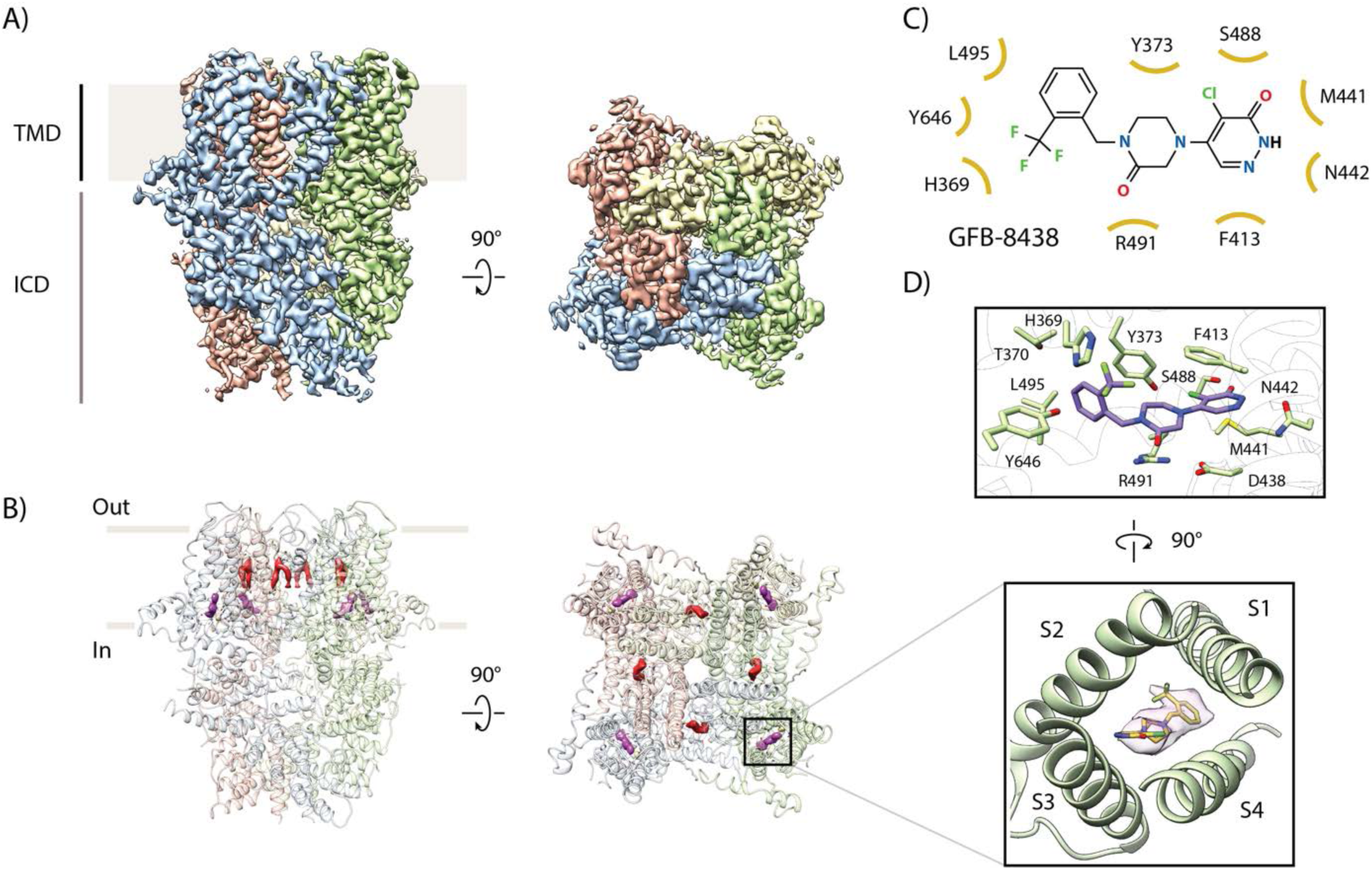
Cryo-EM structure of inhibitor-bound TRPC4 channel. **(A)** Side and top view of the cryo-EM map of GFB-8438 inhibitor-bound TRPC4, with each subunit colored differently. Positions of the transmembrane domain (TMD) and intracellular cytosolic domain (ICD) are indicated. **(B)** Location of non-protein densities relative to the atomic model of TRPC4, which is shown in transparent ribbon representation in the side- and top view. Densities corresponding to lipids are depicted in red, GFB-8438 density is shown in purple. **(C)** Chemical structure of the TRPC4 inhibitor GFB-8438, with important and interacting residues of TRPC4 highlighted. Non-carbon atoms are colored according to element, with halogens in green, nitrogen in blue and oxygen in red. **(D)** Close-up of ligand binding pocket (bottom panel), showing the density corresponding to the inhibitor GFB-8438 in pink with the ligand structure modelled inside. GFB-8434 is enclosed by the four helices S1 to S4, constituting the VSL domain. A rotated view of the ligand binding pocket is shown in the top panel with important and interacting residues highlighted. GFB-8438 is shown in purple.

**Figure 2.**
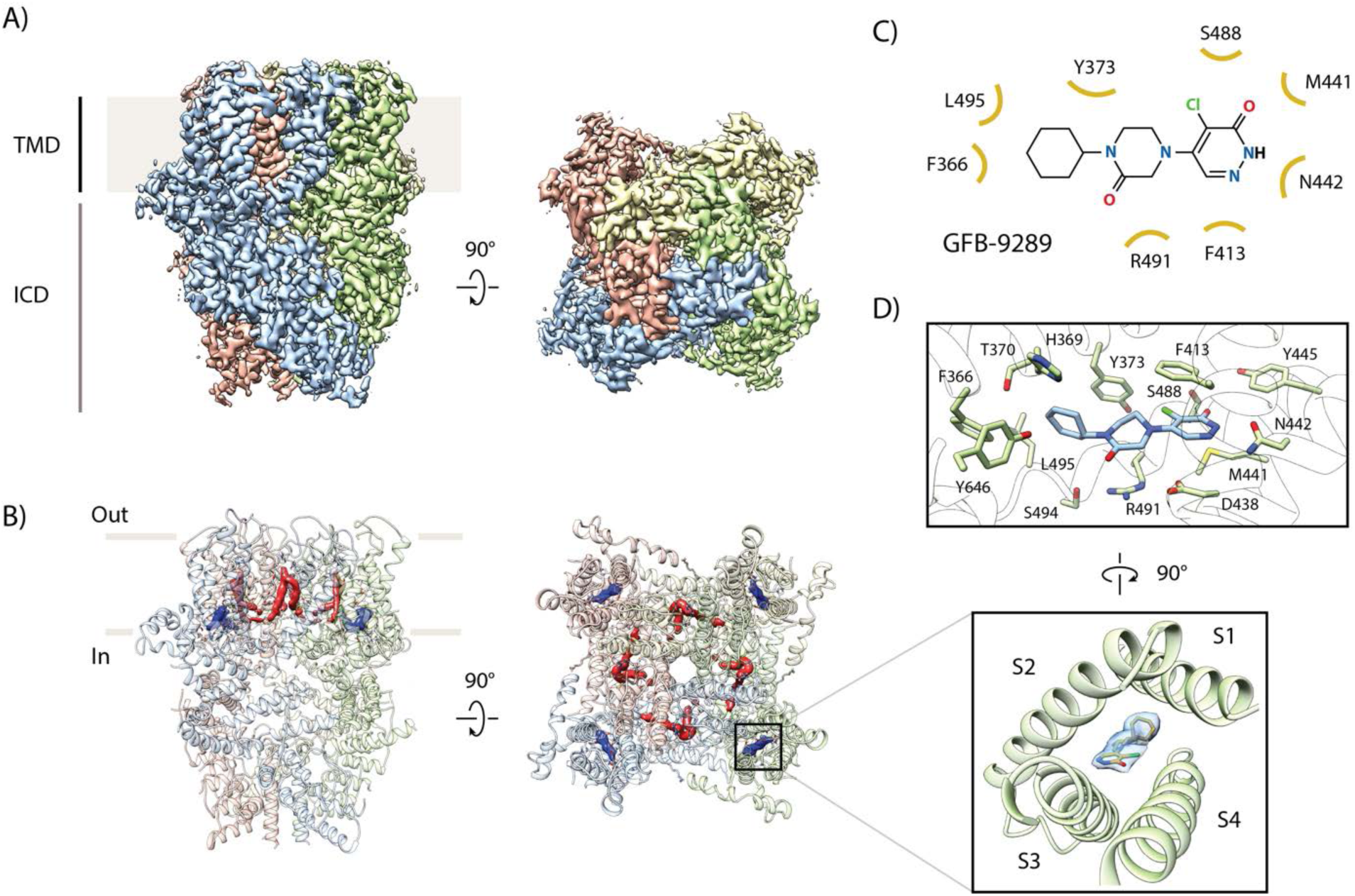
Cryo-EM structure of activator-bound TRPC4 channel. **(A)** Side and top view of the cryo-EM map of GFB-9289 activator-bound TRPC4, with each subunit colored differently. Positions of the transmembrane domain (TMD) and intracellular cytosolic domain (ICD) are indicated. **(B)** Location of non-protein densities relative to the atomic model of TRPC4, which is shown in transparent ribbon representation in the side- and top view. Densities corresponding to lipids are depicted in red, GFB-9289 is shown in blue. **(C)** Chemical structure of the TRPC4 activator GFB-9289, with important and interacting residues of TRPC4 highlighted. Non-carbon atoms are colored as in Figure 1. **(D)** Close-up of the ligand binding pocket (bottom panel), showing the density corresponding to the activator GFB-9289 in blue with the ligand structure modelled inside. The activator is enclosed by the four helices S1 to S4 of the VSL domain. A rotated view of the ligand binding pocket is shown in the top panel with important and interacting residues highlighted. GFB-9289 is shown in blue. Both the activator and inhibitor occupy the same ligand binding pocket.

Overall, the structures of these complexes are similar to the previously determined TRPC4 apo structure (Figure 1A, 2A, Figure S4). The architecture is typical for the canonical TRP channel family with a transmembrane region (TM domain) where the pore region of one protomer domain-swaps with the voltage-sensor-like (VSL) domain of another. The cytosolic domain harboring the ankyrin repeat (AR) embraces the coiled coil helix in the center and the N-terminal ankyrin domain is associated to the TM domain via a helical linker domain. The C-terminal helix connects to the TM domain through the rib and TRP helix which also bridges the TM domain with the helical linker domain (Figure S5).

### TRPC4 in complex with inhibitor GFB-8438

In the GFB-8438-bound structure, we found an additional density compared to the apo structure inside a cavity formed by the VSL domain, TRP helix and re-entrant loop (Figure 1B). The shape of the density clearly indicated that it corresponds to the bound inhibitor. The shape and the surrounding chemical environment allowed us to build the model of the inhibitor inside this extra density (Figure 1C,D). Notably, the inhibitor AM-1473, which belongs to a different class of small molecules, was shown to bind to a similar region in TRPC6 (Bai et al., 2020).

The chemical structure of GFB-8438 consists of three six-membered rings: a pyridazinone ring and trifluoromethyl benzyl group at opposing ends are connected by a central 1,4-disubstituted piperazinone ring (Figure 1C). Its binding to the protein is predominantly mediated by hydrophobic contacts (Figure 1C,D). The nitrogen, the chlorine, and the oxo group of the pyridazinone ring form hydrogen bonds as well as halogen bonds with N442 of helix S3, Y373 of helix S1 and S488 of the S4 helix, respectively. The hydrophobic part of the pyridazinone ring is stabilized by a π-π stacking interaction with F413 of helix S2 on one side and M441 of helix S3 on the opposite side. The middle piperazinone ring forms a hydrophobic interaction with the Y373 while the oxo-group of the ring is engaged in a hydrogen bond with R491 of the S4 helix. The tri-fluoro benzyl ring engages in a hydrophobic interaction with L495 of helix S4, and the fluoride group is involved in a hydrogen bond with H369 of S1 and Y646 from the TRP helix. The residues interacting with the inhibitor are identical between TRPC4 and TRPC5 (Figure S6) indicating a similar ligand binding mode in TRPC5, which is supported by their close IC_50_ values of 0.18 and 0.29 µM for TRPC5 and TRPC4, respectively (Yu et al., 2019).

### TRPC4 in complex with activator GFB-9289

As in the GFB-8438 inhibitor-bound structure, we found an extra density inside the VSL domain region of activator-bound TRPC4 (Figure 2A, B). In addition to the surrounding chemical environment, the high resolution of the map enabled us to unambiguously build the ligand (Figure S3).

Similar to the inhibitor, the chemical structure of the activator consists of three six-membered rings: a pyridazinone ring and a cyclohexyl group at opposing ends are connected by a central 1,4-disubstituted piperazinone ring (Figure 2C). The key difference between the molecules is terminal ring, which is a tri-fluorinated benzyl ring in the case of the inhibitor and a cyclohexyl ring in the activator. Given that the activator and the inhibitor share a common chemical scaffold structure with the difference limited to one part of the molecule, it is not surprising that they bind to the same region. The competitive binding of the compounds to TRPC4 is consistent with observed functional data. After preincubation with the inhibitor, the activator does not have a measurable functional effect on the channel (Figure S1). However, considering the same binding mode and similar structure, it is intriguing that binding of the two related small molecules have opposing effects on the activity of the protein. This is reminiscent of TRPM8, where the activator and inhibitor bind to the same pocket of the VSL domain (Diver et al., 2019; Yin et al., 2018). This suggests that the VSL domain is a highly sensitive regulatory domain that responds to subtle stimuli in its small ligand-binding pocket in order to govern the function of this large tetrameric macromolecular complex. In addition, such opposing effects of closely related small molecules have also been described in the case of TRPC5 (Rubaiy et al., 2018).

Most interactions of the activator with the S1-S4 helices are the same as they are for the inhibitor, with small residue movements to accommodate the slightly different structure of the activator (Figure 2C, D). In the case of the activator, GFB-9289, Y373 forms hydrophobic interactions with the cyclohexyl and piperazinone rings. A reconfiguration of the binding interactions to the pyridazinone ring now includes hydrogen bonds to S488 side chain via its oxygen atom, unlike the halogen bond with Y373 observed in the inhibitor complex described above. The reduced size of the cyclohexyl group results in the reorientation of interacting residues. Importantly, Y646 of the TRP helix and H369 of the S2 helix do not interact with the compound and are rotating away from the interface (Figure 3A). This reduced stabilization of the activator (GFB-9289) is the likely cause for the weaker binding in comparison to the inhibitor (GFB-8438, Figure S1).

**Figure 3.**
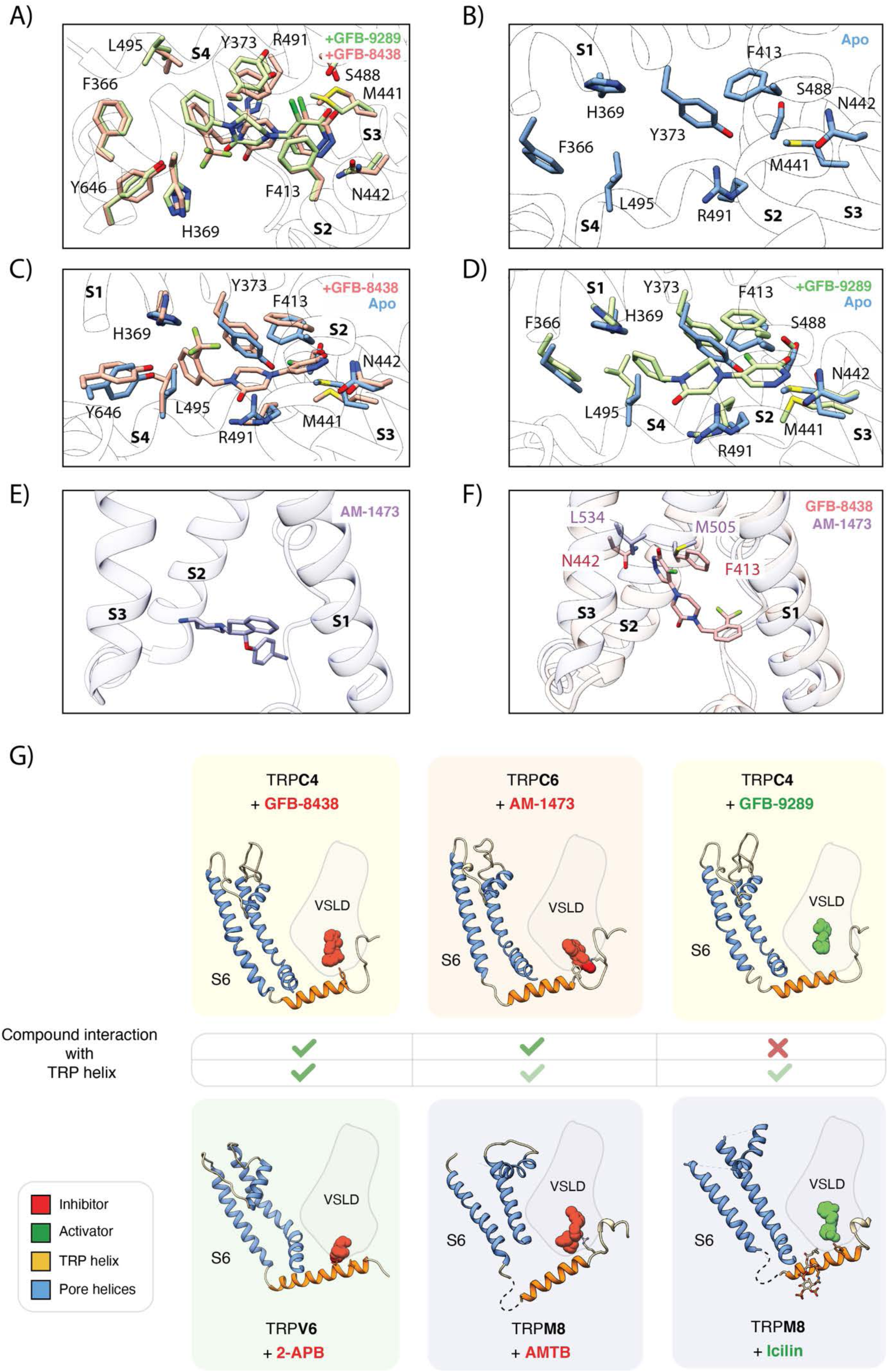
Comparison of the ligand binding pocket in TRPC4. **(A)** Superposition of GFB-8438 inhibitor- (red) and GFB-9289 activator- (green) bound TRPC4 structures. The respective ligand is depicted in the same color as the corresponding residues. Positions of the surrounding helices S1 to S4 are indicated. Due to their structural similarity both molecules adopt a similar orientation within the ligand binding pocket. **(B)** Close-up of ligand binding pocket in the apo TRPC4 structure, which is enclosed by the four helices S1 to S4 of the voltage sensing-like domain. **(C)** Superposition of inhibitor-bound (red) and apo (blue) structure of TRPC4. A close-up of the ligand binding pocket is shown, with important and interacting residues highlighted. The inhibitor GFB-8438 is depicted in red, positions of the surrounding helices S1 to S4 are indicated. **(D)** Same as in (C) for the activator-bound TRPC4 structure, which is shown in green. The structure of the activator GFB-9289 is also depicted in green. In both the activator- and inhibitor-bound structures, several residues move away from the center of the pocket to create space for accommodating the respective ligand. **(E)** Position of the inhibitor AM-1473 within the VSL domain binding pocket of TRPC6 is shown. The surrounding helices S1-S3 are indicated for orientation. **(F)** Superposition of GFB-8438-bound TRPC4 (red) and AM-1473-bound TRPC6 (purple) channels. The location of the GFB-8438 inhibitor within the VSL domain is shown. In contrast to AM-1473, which is located in the lower part of the binding pocket (see E), GFB-8438 additionally interacts with the upper region of the pocket. The depicted residues in this region contribute to the selectivity of GFB-8438 for TRP4/5 channels. **(G)** Comparison of small molecule modulators of the TRP channel family that target the ligand binding pocket enclosed by the helices of the VSL domain (VSLD). Small molecules are depicted as space-filled spheres and colored according to their respective modulatory effect, with inhibitors in red and activators in green. Residues interacting with the ligand are shown in stick representation. Pore-forming helices are colored in blue, the TRP helix in orange. Within the TRPC subfamily, inhibitors interact with the TRP helix, whereas activators do not.

### Structural rearrangements in the ligand binding pocket

To understand the structural rearrangement upon ligand binding, we compared the ligand-bound structures with the structure of TRPC4 in the apo state (Vinayagam et al., 2018) (Figure 3B-D). In the apo structure, some of the residues of the ligand binding pocket interact with each other via hydrophobic (Y373, F413, M441) and hydrophilic interactions (R491 and E438) (Figure 3B). Upon ligand binding, these residues move and reshape the pocket to accommodate the ligands, indicating an induced-fit mechanism or conformational selection (Hammes et al., 2009) (Figure 3B-D). Similarly, the side chains of L495 and H369 rotate, move or flip to accommodate and stabilize the interaction with the tri-fluoro benzyl group in the case of the inhibitor and arrange differently in the case of the activator (Figure 3B,C). These ligand-specific arrangements of the ligand binding pocket highlight its plasticity.

The inhibitor GFB-8438 has been shown to be more specific for TRPC4/5 than TRPC6 (Yu et al., 2019). Comparison of the TRPC4/5 binding pocket with TRPC6 reveals a critical difference in ligand binding residues (Figure 3D). The cognate N442 residue in TRPC4/5 is replaced by L534 in TRPC6, which abrogates hydrogen bond formation with the nitrogen atom of pyridazinone. F413, which in TRPC4 forms a π-π interaction with the pyridazinone ring, is replaced by the weakly interacting hydrophobic residue M505 in TRPC6 (Figure 3E,F). These crucial substitutions in the ligand binding pocket explain the much lower binding affinity of TRPC6 for GFB-8438 (>30 μM) (Yu et al., 2019).

Interestingly, we have observed that the inhibitor, GFB-8438, interacts with both the TRP helix and the VSL domain, whereas the activator, GFB-9289, exclusively interacts with the VSL domain (Figure 3G). We hypothesize that the direct stabilizing interaction with the TRP helix constrains it and the adjacent S6 helix thereby arresting the channel in a closed state. In this manner, the allosteric interactions within a peripheral binding site propagate to the ion pore in the center of the protein. To explore our hypothesis further, we compared the inhibitor binding to the VSL domain observed by us with that of other TRP family members. Indeed, all inhibitors interact with the TRP helix. This indicates that the above described mechanism may also be generally valid for other inhibitors that bind to the VSL domain of different TRP channels. However, in the case of TRPM8, both the activator and inhibitor were found to interact with the TRP helix. Yet the connecting loop between the TRP helix to the S6 helix was disordered in both structures. Therefore, we believe that TRPM8 utilizes a different mechanism to conduct the ligand mediated signal than the TRP channels examined here (Figure 3G).

### Ligand-induced changes in TRPC4

Besides the structural rearrangements in the ligand binding pockets, we did not observe major ligand-induced conformational changes in TRPC4. Similar to the apo structure, the channel is closed at the lower gate in the structure of the inhibitor-bound TRPC4. The lower gate shows a minimal constriction defined by residue N621 with a van der Waals surface diameter of approximately 0.7 Å, which is too narrow for Ca^2+^ to pass through (Figure 4A-B). Surprisingly, the activator-bound structure shows also a closed channel. The recently reported structure of TRPC6 bound to an activator also exhibits a closed conformation (Bai et al., 2020). The opening probability of TRPC channels is generally very low and the opening time is short (<1 ms) (Hofmann et al., 1999; Jung et al., 2003; Schaefer et al., 2000). Therefore, even in the presence of an activator, the closed state seems to be energetically more favorable than the open state.

**Figure 4.**
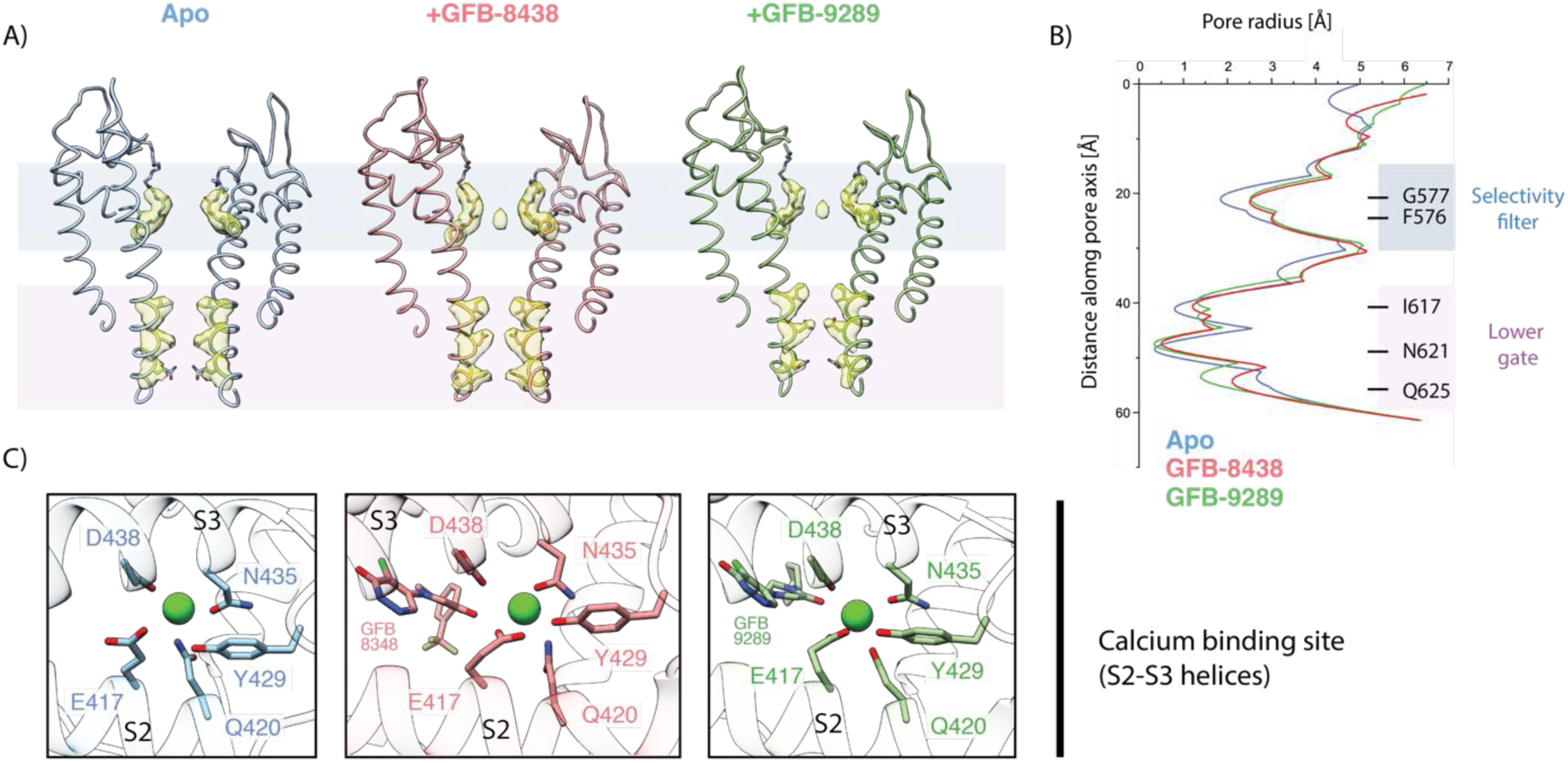
Comparison of the ion conduction pore and Ca^2+^-binding site. **(A)** Side view of the pore-forming region of TRPC4 in the apo- (blue), GFB-8438 inhibitor-bound (red) and GFB-9289 activator-bound (green) structures. Only the two opposing subunits of the tetrameric channel are shown as ribbon representation for clarity. The density at comparable thresholds corresponding to the selectivity filter (light blue) and the lower gate (pink) is superimposed. A central density is observed in all maps, except the apo structure. **(B)** The calculated pore-radii corresponding to the three TRPC structures in (A) are depicted. The color code is also identical to (A). The positions of important residues, constituting the selectivity filter and the lower gate, are indicated on the right. **(C)** Close-up of the Ca^2+^-binding site in the three TRPC4 structures, located in direct vicinity to the ligand binding pocket of the VSL domain. Position of ligands and coordinating residues are highlighted. Color code of TRPC4 structures is as in (A).

We found small differences in the selectivity filter. Surprisingly, the backbone residues F576 and G577 forming the TRPC4 selectivity filter show a slightly wider radius in the ligand-bound structures compared to the apo structure. This could be due to a density in the selectivity filter which we did not observe in the apo structure, indicating that a cation, presumably Ca^2+^ or Na^+^ is residing in the filter, while the filter is empty in the apo form (Figure 4A).

In both ligand-bound structures a characteristic lipid density is situated close to the pore which is either phosphatidic acid or ceramide-1-phosphate, its structural analogue (Vinayagam et al., 2018). Since we did not add lipids during our purification, this annular lipid likely co-purified with the protein. Each of the two lipid tails is placed like an anchor between neighbouring S5 and S6 helices by forming several hydrophobic interactions (Figure S7). We hypothesize that this lipid site could be crucial for the gating of the channel, since small molecules can bind in this region, and modulate the channel as observed in activator-bound TRPC6 (Bai et al., 2020).

We identified density for a putative cation in the Ca^2+^ binding site of the VSL domain in the ligand-bound structures (Figure 4C and S8A), also present in the apo structure of TRPC4 (Vinayagam et al., 2018). The ion binding site is coordinated by the carbonyl oxygen of D438 and N435 of the S3 helix, along with E417 and Q420 of the S2 helix, which are the favourable coordination residues for an alkaline earth metal ion such as Ca^2+^ (Zheng et al., 2017a). Interestingly, not only a hydroxyl group of Y429 is close to the density but also an oxo group of the ligands, which could complete the octahedral coordination of Ca^2+^ via a bridging water molecule. The presence of the ligands could thus help to stabilize bound Ca^2+^.

TRPC5 and TRPC4 activation has been reported to be Ca^2+^-dependent (Plant and Schaefer, 2003). Similar to our observation here, the binding of Ca^2+^ has been described for TRPM4 and TRMP8, both of which are also known to be activated by Ca^2+^. The structures of these channels are in a closed conformation representing the desensitized state (Autzen et al., 2018; Diver et al., 2019). Considering this, the molecular role of the VSL domain-bound calcium ion in activation or desensitization of the TRPC4 channel is a compelling topic for further investigation.

### Structure of TRPC4 in complex with calmodulin

Calmodulin (CaM) has been shown to bind and regulate the TRPC4 channel (Zhu, 2005; Tang et al., 2001). At high Ca^2+^ concentrations in the cytosol, CaM binds in its Ca^2+^-bound state to TRPC4 and inhibits Ca^2+^ entry. At low Ca^2+^ concentrations, CaM changes its conformation and dissociates from the channel. The store-operated Ca^2+^ entry pathway hypothesis (Tang et al., 2001) further proposes that CaM binding to the channel at resting state prevents TRPC4 from being spontaneously activated by IP_3_ receptors. When Ca^2+^ levels in the endoplasmic reticulum (ER) - but not in the cytosol - drop, the affinity of the IP_3_ receptor to TRPC4 increases and CaM is displaced through a conformational coupling mechanism (Rosado et al., 2015; Tang et al., 2001). This activates the TRP channel. To further understand the mechanistic process of CaM inhibition, we set out to determine the structure of the TRPC4-CaM complex.

We first performed a pull-down experiment using a CaM Sepharose column with TRPC4 acting as bait at high Ca^2+^ concentrations to biochemically test CaM binding to TRPC4. As expected, TRPC4 was trapped in the CaM column in presence of Ca^2+^ and released by chelating the Ca^2+^ with EGTA (Figure S9). This is in line with previous studies which used smaller peptides of TRPC4 instead of the full-length protein used in our experiment (Tang et al., 2001). Since Ca^2+^ is necessary for the binding of CaM to TRPC4, we prepared the protein sample in the detergent LMNG (Lauryl Maltose Neopentyl Glycol) instead of following the amphipol exchange methodology that we used previously. Amphipols are known to interact with Ca^2+^ ions and could thus disrupt CaM binding (Le Bon et al., 2018). We then determined the structure of TRPC4 in LMNG in complex with CaM. For the TRPC4-CaM complex, we added a 10-fold molar excess of CaM to tetrameric TRPC4 in the presence of 10 mM calcium chloride throughout the purification process after detergent extraction.

The CaM complex sample yielded a 3.3 Å map with applied C4 symmetry (Figure S10). We observed additional density surrounding the rib helix termini protruding from the protein core, although the resolution in this region was lower than at the core of the protein (Figure S11a). Besides its localization at the periphery we suspected that an incomplete saturation of TRPC4 by CaM could be the reason for the lower local resolution. Hence, we performed 3D sorting without applied symmetry to resolve the subpopulations with different binding stoichiometries. 13% of the TRPC4 channels had one CaM bound, 35% and 31% had two or three bound, respectively and only 20% were fully saturated (Figure 5A). In addition, some of the densities corresponding to CaM were less defined than others. The classes with clear CaM densities were then rotated and properly aligned (Figure S10). The final local resolution of CaM improved to a resolution of 4 - 5 Å (Figure S11a). We could clearly identify four helices that correspond to the helices of one lobe of CaM and flexibly fitted this part of the protein (Figure 5B). The other CaM lobe was not resolved, indicating that this part of the protein is more flexible in this complex.

**Figure 5.**
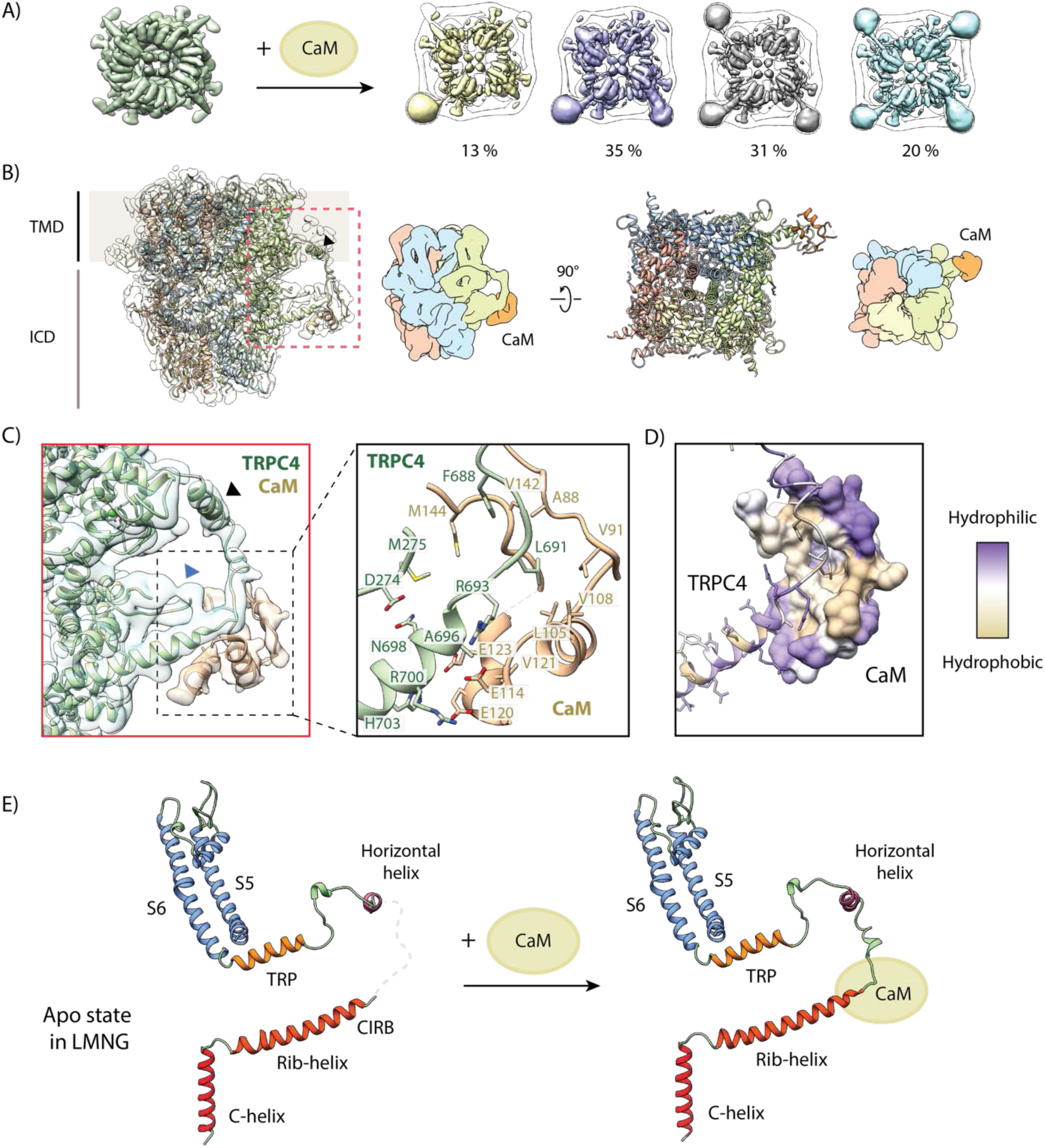
Structural basis for inhibition of TRPC4 by calmodulin. **(A)** One to four CaM molecules are bound to the CIRB binding sites of the tetrameric TRPC4 channel. 13% of particles are decorated with one (yellow), 35% with two (lilac), 31% with three (grey) and 20% with four CaM molecules (turquoise). **(B)** Side view of the CaM-bound TRPC4 density map (transparent) with the corresponding atomic model fitted inside, in which each protomer is colored differently. Position of the horizontal helix is indicated by black arrowhead. The bottom view of the atomic model is shown in the right panel. A schematic representation for both views is provided next to the pdb models. CaM is colored in orange. **(C)** Close-up of the indicated region in (B), showing the CaM binding region (left panel). CaM is colored in orange, TRPC4 in green. Positions of the horizontal helix and loop region 273-277 are indicated by black and blue arrowhead, respectively. Important and interacting residues of TRPC4 and CaM are highlighted in the right panel. **(D)** TRPC4 (cartoon representation) and CaM (surface representation) are colored according to hydrophobicity. There is central hydrophobic cavity in CaM that is surrounded by hydrophilic residues in its periphery. The complementary binding region of TRPC matches this profile. **(E)** The C-terminal helix (red), the rib-helix (red-orange), the horizontal helix (purple), the TRP helix (orange) and the pore-forming helices (blue) of a single TRPC4 promoter are shown before (left panel) and after CaM binding (right panel). CaM binding stabilizes the previously disordered region connecting the rib-helix and horizontal/TRP-helix. LMNG – lauryl maltose neopentyl glycol

CaM not only binds to the tip of the rib helix (residues 691-703) and the following loop (residues 677-690) that connects the rib helix with a newly identified helix (residues 666-676), but it also interacts with the adjoining loop region comprising residues 273-277 (Figure 5C). The core region of CaM binds to TRPC4 by forming hydrophobic interactions while the peripheral residues of CaM are stabilized by hydrophilic interactions (Figure 5D) that are typically observed in CaM-protein/peptide complexes (Villalobo et al., 2018).

The interacting residues of TRPC4 partially overlap with a peptide corresponding to residues 695-724 that have been previously shown to interact with CaM (Tang et al., 2001). Since our structure revealed that CaM only interacts with residues 688-703, we conclude that the residues 695-703 are sufficient for CaM binding *in vitro*. Residues 704-725 of the rib helix interact with the protein core and are inaccessible for interaction with CaM.

### CaM-induced changes in TRPC4

To be able to identify CaM-induced structural effects, we also solved the structure of TRPC4 in its apo state under the same conditions as for the TRPC4-CaM complex without the addition of external lipids. The apo structure of TRPC4 in LMNG reached a resolution of 2.85 Å, allowing us to build an atomic model with high accuracy (Figure S11B and S12). The overall structure is similar to the previously reported amphipol-exchanged apo structure of TRPC4 in the closed state (Vinayagam et al., 2018) (Figure S4). However, in both the apo and calmodulin-bound structure, we observed for the first time an additional density corresponding to a horizontal helix located at the transmembrane-cytoplasmic interface outside the transmembrane core (residues 666-676) (Figure 5B-D, Figure S12). The hydrophobic residues of this helix face the transmembrane helix and the inner lipid leaflet while the hydrophilic residues project into the cytoplasm, giving the helix an amphipathic nature.

Comparing the TRPC4 apo structure with that of TRPC4-CaM, we could identify only small differences in the center of the channel. Both, the apo and CaM bound TRPC4 structures showed the same constriction of 0.7 Å defined by N621 at the lower gate indicating the closed state of the channel (Figure S13). Interestingly, when CaM binds to TRPC4 the selectivity filter is slightly widened and contains a density that likely corresponds to Ca^2+^ or Na^+^ as in the case of the activator- and inhibitor-bound state. We observed a much stronger density at the Ca^2+^ binding site in the VSL domain for the CaM-bound structure compared to apo TRPC4, presumably due to the high Ca^2+^ concentration we used for preparing the CaM-TRPC4 complex The differences between TRPC4-CaM and the TRPC4 apo structure are more pronounced at the periphery of the channel. There, CaM binding stabilizes a longer stretch (residues 677 −692) of TRPC4 that is highly flexible in the apo state of the channel (Figure 5B and C). Therefore, binding of CaM to this region of TRPC4 likely reduces the overall flexibility of the channel, fixing it in its closed state (Figure 5E). Since one TRPC4 tetramer can bind up to four CaMs, this suggests that the number of CaMs simultaneously bound to TRPC4 could fine tune the level of channel activity.

Importantly, this mechanism of CaM-mediated regulation completely differs from that described for other TRP channels, such as TRPV5 and TRPV6 (Hughes et al., 2018; Singh et al., 2018). There, CaM binds in a 1:4 stoichiometry, with one CaM binding to the center of the tetrameric channels via the open cytoplasmic part, plugging it with its protruding lysine residue (Figure 6) (Hughes et al., 2018; Singh et al., 2018). In the TRPC4-CaM complex structure, the central core of the cytoplasmic region is occupied by a coiled coil helix. Thus, CaM cannot access the core of the cytoplasmic region in TRPC4. Other channels of the TRPC subfamily also contain this coiled coil helix and the rib helix (Duan et al., 2018; Tang et al., 2018). Therefore, we propose that the novel mechanism of CaM inhibition via binding to the rib helix is paradigmatic for all TRPCs.

**Figure 6.**
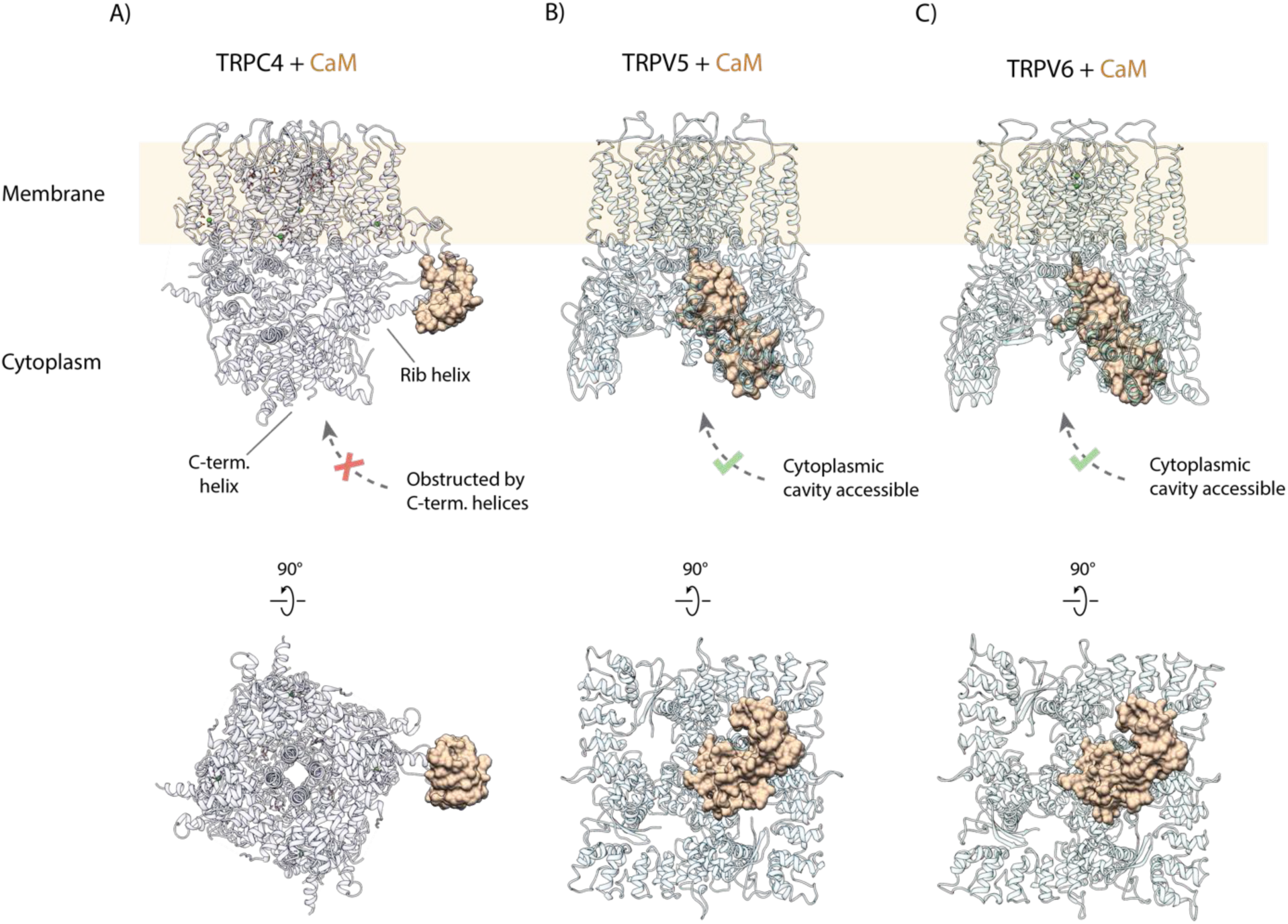
Comparison of CaM binding in TRPC and TRPV channels. **(A)** Calmodulin (CaM) interacts with the rib helix of TRPC4. Side (upper panel) and bottom (lower panel) view of the CaM-bound TRPC4 is shown, with TRPC4 structure in cartoon representation with moderate transparency and CaM in space filling sphere representation. Only a single lobe of the double-lobed CaM molecule is resolved in the structure. This indicates that the second lobe is rather flexible. Up to four binding sites are accessible for CaM (only one binding event is shown here for clarity). **(B)** Same as in (A) for TRPV5. The two-lobed CaM binds into the central cytoplasmic cavity of TRPV5. While four potential binding sites are available in TRPV5, only a single CaM molecule can bind due to steric hindrance. Unlike TRPC4, in which the C-terminal helices block the access to the cytoplasmic cavity, CaM can enter into the internal cavity of TRPV5 from the cytoplasm. **(C)** Same as in (A) for TRPV6. Similar to TRPV5, only a single CaM molecule binds to a region within the cytoplasmic cavity of TRPV6, indicating that this binding mode is conserved among TRPV channels.

### Model for TRPC4 modulation

In this study we determined the structure of TRPC4 in complex with inhibitor GFB-8438, activator GFB-9289 and its endogenous regulator CaM. Analysis of these structures allows us to propose a model describing the molecular mechanism of modulation and regulation of TRPC4 activity (Figure 7a). In our model, the channel switches between its closed and open conformation, with the closed conformation being more energetically favoured. Therefore, activation does not result in a static opening of the channel, but rather increases the open time interval. The channel only transiently opens and allows the passage of Ca^2+^. Therefore, we also only obtained the structure of TRPC4 in its closed conformation although the activator is bound to the protein. Inhibition has the reverse effect, i.e. the open time interval is strongly decreased and an inhibitor locks the channel in its closed conformation. In our case, the inhibitor and the activator bind to the same position, namely the VSL domain which is connected to the gate by the TRP helix. Thus, subtle conformational changes in this sensitive regulatory domain appear to shift the equilibrium between the open and closed states.

**Figure 7.**
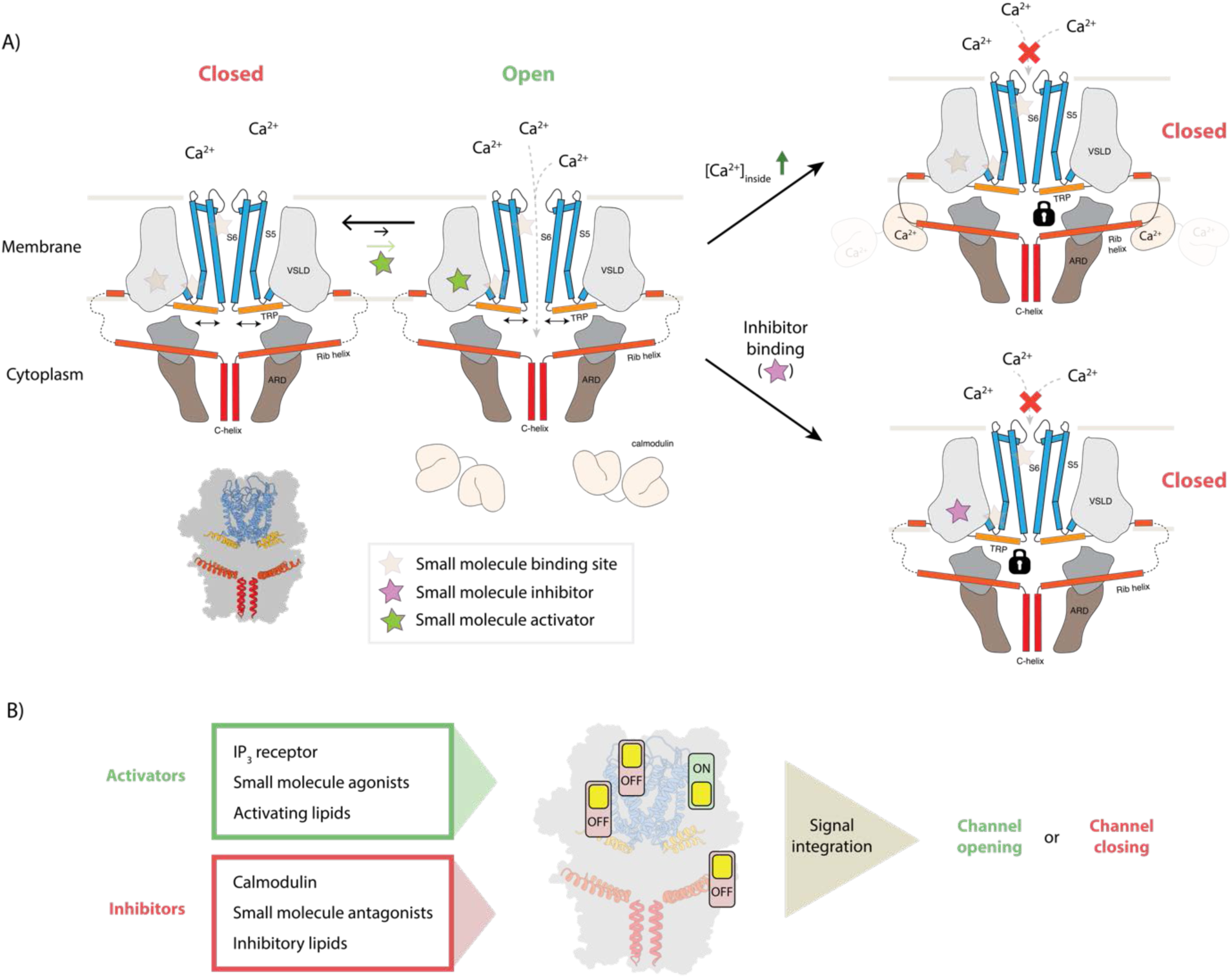
Model for TRPC4 modulation. **(A)** Canonical TRP channels can transiently open to allow the passage of Ca^2+^ ions into the interior compartment (left panel). Several mechanisms modulate the activity of the channel: binding of small molecule activators to one of the ligand binding pockets favors the opening event and thereby increases the overall channel activity. In the gating process, the TRP helix (orange) plays a central role as it has a direct connection to the pore-forming helices (blue), constituting the ion conducting pore. Binding of small molecule inhibitors and the inhibitory protein CaM can restrict the mobility of the TRP helix, thus locking the channel in the closed state (bottom and top panels on the right, respectively). In the latter case, high intracellular Ca^2+^ concentrations cause the Ca^2+^-sensing protein CaM to bind to the CIRB region of the protruding rib-helix (red). This binding event stabilizes a previously disordered region that directly connects to the TRP helix. **(B)** Individual or simultaneous binding of activators and/or inhibitors modulate the channel gating. Interestingly, modulation sites, i.e. ligand pockets or structural features to which certain compounds or regulatory proteins bind, can accommodate both activators and inhibitors. Thus, these regions can be considered as activity switches. Binding of activators results in an “ON” position, whereas inhibitor binding causes an “OFF” state. In the case that multiple modulators bind simultaneously, all signals are integrated to determine whether the channel opens or remains closed.

CaM does not bind to the VSL domain, which resides in the membrane and is therefore not directly accessible. However, it stabilizes other parts of the protein that are connected to the VSL domain. In particular, it binds to the tip of the rib helix, which results in the stabilization of the loop and the helix that connects it to the VSL domain. Thus, the binding of CaM to the rib helix has the same consequence as the binding of an inhibitor to the VSL domain, locking the channel in its closed conformation. Interestingly, the rib helix has also been shown to be the binding site for the IP_3_ receptor which acts as an activator of TRPC4 (Tang et al., 2001). Although we do not yet know the structural details of this interaction, it is likely that inhibition by CaM and activation by IP_3_ receptor require the use the same binding site, while resulting in opposing effects. This is similar to the activator/inhibitor pair binding to the VSL domain. Thus, TRPC4 contains at least two molecular switch regions that can be modulated by the binding of small molecules or regulatory proteins (Figure 7b). Consequently, the signals of different switches are integrated and together determine the final state and the degree of activation of the channel. Our model not only explains how TRPC4 activity is regulated by CaM in the cellular context, but also opens new possibilities for knowledge-driven pharmacological manipulation of this therapeutic target.

## Materials and Methods

### Protein purification and expression

Zebrafish TRPC4_DR_ was prepared as described previously (Vinayagam et al., 2018). In brief, residues 2-915 of *Danio rerio* TRPC4 were cloned into the pEG BacMam vector (Goehring et al., 2014), with a C-terminal HRV-3C cleavage site followed by EGFP, and a twin StrepII-tag. An 8x His-tag with a TEV cleavage site was positioned at the N-terminus. Baculovirus was produced as described previously (Goehring et al., 2014). The P2 baculovirus produced in Sf9 cells was added to HEK293 GnTI^-^ cells (mycoplasma test negative, ATCC #CRL-3022) and grown in suspension in FreeStyle medium (GIBCO-Life Technologies) supplemented with 2% FBS at 37°C and 8% CO_2_. After 8 hours of transduction 5 mM sodium butyrate was added to enhance protein expression and allowed the cells to grow for an additional 40 hours at 30°C.

48 hours post transduction, cells were harvested by centrifugation at 1,500 *g* for 10 mins and washed in phosphate-buffered saline (PBS) pH 7.4. The cell pellet was resuspended and cells were lysed in an ice-cooled microfluidizer in buffer A (PBS buffer pH 7.4, 1 mM Tris(2-carboxyethyl) phosphine (TCEP), 10% glycerol) in the presence of protease inhibitors (0.2 mM AEBSF, 10 µM leupeptin). 50 ml buffer A was used per pellet obtained from 800 ml of HEK293 cell culture. The lysate was centrifuged at 5,000 *g* for 5 min to remove cell debris, followed by a 15,000 g centrifugation for 10 mins to remove sub-cellular organelles. The membranes were collected by ultracentrifugation using a Beckman Coulter Type 70 Ti rotor at 40,000 rpm. The membranes were then mechanically homogenized in buffer B (100 mM Tris-HCl pH 8, 150 mM NaCl, 1 mM TCEP, 10% glycerol) containing protease inhibitors, flash-frozen and stored at −80 °C until further purification.

### Purification of TRPC4 in DDM followed by amphipol exchange

Membranes were solubilized for 2 hr in buffer B supplemented with 1% dodecyl maltoside (Anatrace #D310). Insoluble material was removed by ultracentrifugation for 1 hr in a Beckman Coulter Type 70 Ti rotor at 40,000 rpm. The soluble membrane fraction was diluted 2-fold with buffer B and applied to a column packed with Strep-Tactin beads (IBA Lifesciences) by gravity flow (6–10 s/drop) at 4 °C. Next, the resin was washed with ten column volumes of buffer B supplemented with 0.04% DDM solution containing protease inhibitors. Bound protein was eluted seven times with 0.5 column volumes of buffer A with 3 mM D-Desthiobiotin (Sigma-Aldrich), 0.026% DDM and 0.1 mM AEBSF protease inhibitor. The C-terminal EGFP tag was removed by incubating the eluted fractions with HRV-3C protease overnight. The next day, the detergent was replaced with amphipols A8-35 (Anatrace) 4:1 (w/w) to the cleaved protein and incubating for 6 hr at 4 °C. Detergent removal was performed by adding Biobeads SM2 (BioRad) pre-equilibrated in PBS to the protein solution at 10 mg/ml final concentration for 1 hr, then replaced with fresh Biobeads at 10 mg/ml for overnight incubation at 4 °C. Biobeads were removed using a Poly-Prep column (BioRad) and the solution was centrifuged at 20,000 g for 10 min to remove any precipitate. The protein was concentrated with a 100 MWCO Amicon centrifugal filter unit (Millipore) and purified by size exclusion chromatography using a Superose 6 Increase 10/300 GL column (GE Healthcare) equilibrated in buffer C (PBS pH 7.4, 1 mM TCEP). The peak corresponding to tetrameric TRPC4_DR_ in amphipols was collected and analysed initially with negative stain EM and then by cryo-EM.

### TRPC4 pulldown assay using CaM sepharose beads

The assay was performed with manufacturer instructions using TRPC4 as a bait. Briefly, 1 ml of CaM sepharose beads were loaded into the Biorad Ployprep column and washed with 10 ml of binding buffer containing 20 mM Tris-HCl (pH 7.5), 150 mM NaCl, 2 mM CaCl_2_. TRPC4 prepared in LMNG (described below) was loaded onto the column by gravity flow (10 s/drop) at 4 °C. After loading, the column was washed with 10 ml of binding buffer. Finally, TRPC4 was eluted with 5ml of elution buffer containing 20 mM Tris-HCl (pH 7.5), 150 mM NaCl, 2 mM EGTA (Figure S9).

### Purification of CaM

Mouse CaM was subcloned into a pET19 vector and expressed in BL21-CodonPlus (DE3) - RIPL cells. Cells were grown in LB broth with 125 µg/ml ampicillin at 37 °C until an OD600 of 0.4 was reached. Subsequently, CaM expression was induced with 1 mM IPTG and grown overnight at 20 °C. Cells were harvested by centrifugation and resuspended in 50 ml (per liter of culture) of lysis buffer containing 20 mM Tris-HCl (pH 8.0), 150 mM NaCl and 5 mM imidazole. The cells were lysed in an ice-cooled microfluidizer. The soluble fraction obtained after centrifugation was loaded onto an 8 ml Talon resin column pre-equilibrated with lysis buffer. The resin was washed with 100 ml of lysis buffer containing 20 mM Tris-HCl (pH 8.0), 150 mM NaCl and 20 mM imidazole before eluting in 5 x 5 ml fractions using 25 ml of lysis buffer supplemented with 20 mM Tris-HCl (pH 8.0), 150 mM NaCl and 250 mM imidazole. Calmodulin was further purified by size exclusion chromatography using a Superose 6 10/300 gel filtration column and stored at −80 °C in a storage buffer consisting of 20 mM Tris-HCl (pH 8.0), 150 mM NaCl, 10 % glycerol.

### Preparation of the TRPC4-CaM complex

TRPC4 membranes were solubilized for 2 hr in buffer B supplemented with 1% LMNG (Anatrace #NG310). Then a protocol similar to that used for DDM purification was followed, except that DDM in buffer B was replaced by LMNG with the addition of 10 µM calmodulin and 10 mM calcium chloride. The LMNG detergent concentration was maintained at 5 times the CMC for washing buffer and 3 times CMC for elution. The C-terminal EGFP tag was removed by incubating the eluted fractions with HRV-3C protease overnight. The complex was further purified by size exclusion chromatography using a Superose 6 Increase 10/300 GL column (GE Healthcare) equilibrated in buffer containing 20 mM Tris-HCl (pH 8.0), 150 mM NaCl, 1 mM TCEP, 10 mM calcium chloride and 5% glycerol. Complex formation was assessed by running SDS-PAGE of the peak fraction known to contain TRPC4 (Figure S9). The gel analysis indicated sub-saturation of the complex. Hence, 10 µM CaM was added to saturate the complex before concentrating it to 0.3 mg/ml for plunging. The preparation of TRPC4 -apo in LMNG was similar to the TRPC4-CaM complex except that CaM and CaCl_2_ were not added.

### Cryo-EM grid preparation and screening

The sample quality and integrity were evaluated by negative stain electron microscopy prior to cryo-EM grid preparation and image acquisition as described earlier (Vinayagam et al., 2018). Typically, 4 µl of TRPC4_DR_ at a sample concentration of 0.02 mg/ml were applied onto a freshly glow-discharged copper grid with an additional thin carbon layer. After incubation for 45 s, the sample was blotted with Whatman no. 4 filter paper and stained with 0.75% uranyl formate. The images were recorded manually with a JEOL JEM-1400 TEM operated at an acceleration voltage of 120 kV, and a 4k F416 CMOS detector (TVIPS). For cryo-EM the ligands dissolved in DMSO were added to a final concentration of 100 µM (final DMSO concentration 1%) to TRPC4 exchanged in amphipols and incubated for 30 minutes before plunging using a Vitrobot cryo-plunger (FEI Thermo Fisher) operated at 4 °C and 100% humidity. Details of the plunging conditions are summarized in Table 1.

**Table 1.** Plunging and imaging conditions used for cryo-EM analysis of TRPC4 bound with ligands.

| Sample | Grid type | Volume | Concentration | Blotting time | Blotting force |
| --- | --- | --- | --- | --- | --- |
| TRPC4-8438 | C-Flat 2/1 | 3 μl | 0.3 mg/ml | 3 s | -10 |
| TRPC4-9289 | C-Flat 1.2/1.3 | 3 μl | 0.35 mg/ml | 3 s | 0 |
| TRPC4-CaM | QF 2/1 | 3 μl | 0.3 mg/ml | 3 s | 0 |
| TRPC4-apo(LMNG) | C-Flat 1.2/1.3 | 3 μl | 0.4 mg/ml | 3 s | -3 |

1.2 Imaging Conditions
| Microscopy | TRPC4-apo | TRPC4-CaM | TRPC4-GFB9289<br>(activator) | TRPC4-GFB8438<br>(inhibitor) |
| --- | --- | --- | --- | --- |

| Microscope | Titan Krios (X-FEG, Cs-corrected) |  | Titan Krios (X-FEG, Cs 2.7 mm) |  |
| --- | --- | --- | --- | --- |
| Voltage [kV] | 300 |  | 300 |  |
| Defocus range [μm] | 0.65 to 3.02 | 0.38 to 3.48 | 0.68 to 3.64 | 0.35 to 3.52 |
| Camera | K2 counting | K2 counting | K3 Super resolution | K3 Super resolution |
| Pixel size [$\text{\AA}$] | 0.85 | 0.85 | 0.455 /0.91<sup>a</sup> | 0.455/0.91<sup>a</sup> |
| Total electron dose [$\text{e}/\text{\AA}^2$] | 88.7 | 88.2 | 65.45 | 66.58 |
| Exposure time [s] | 10 | 10 | 3 | 3 |
| Frames per movie | 50 | 80 | 60 | 60 |
| Number of images | 2755<br>(3079) | 6937<br>(7972) | 2369<br>(2970) | 4444<br>(4676) |

<sup>a</sup> Pixel size after 2x binning used for processing In parenthesis is the initial number of images.

### Cryo-EM data acquisition and image processing

Data sets were collected using EPU software on Titan Krios microscopes (FEI Thermo Fisher) operated at 300 kV and equipped with an X-FEG. For the dataset of the activator-bound TRPC4 the aberration-free image shift (AFIS) feature of EPU was used to speed up the data-collection process. Equally dosed frames were collected using a K2 Summit (Gatan) or K3 (Gatan) direct electron detectors in super-resolution mode in combination with a GIF quantum-energy filter set to a filter width of 20 eV. The details of all four data sets including pixel size, electron dose, exposure time, number of frames and defocus range are summarized in Table 1. Data collection was monitored live using TranSPHIRE (Stabrin, 2020), allowing for direct adjustments of data acquisition settings when necessary, i.e. defocus range or astigmatism. The total number of images collected is summarized in Table 1. Preprocessing included drift correction with MotionCor2 (Zheng et al., 2017b), creating aligned full-dose and dose-weighted micrographs. The super-resolution images were binned twice after motion correction to speed up further processing steps. CTF estimation was also performed within TranSPHIRE using CTFFIND 4.1.10 (Rohou and Grigorieff, 2015) on non-dose weighted aligned micrographs. Unaligned frame averages were manually inspected and removed based on ice and image quality, resulting in a removal of 5-20 % of the data sets (see Table 1 for details). Following processing steps were performed using motion-corrected dose weighted sums in the SPHIRE software package unless otherwise indicated (Moriya et al., 2017).

Single particles were picked automatically with crYOLO using the general model (Wagner et al., 2019). The particles were then windowed to a final box size of 288 × 288 pixels. Reference-free 2-D classification and cleaning of the data set was performed with the iterative stable alignment and clustering approach ISAC (Yang et al., 2012) in SPHIRE. ISAC was performed at a pixel size of 3.52 Å/pixel for apo-TRPC4 and TRPC4 bound to CaM and the inhibitor, and with 3.8 Å/pixel for the TRPC4-activator complex. The ‘Beautify’ tool of SPHIRE was then applied to obtain refined and sharpened 2-D class averages at the original pixel size, showing high-resolution features. A subset of particles producing 2-D class averages and reconstructions with high-resolution features were then selected for further structure refinement. The previously reported apo structure was used as reference for 3D refinement in MERIDIEN with imposed C4 symmetry (Moriya et al., 2017). Further polishing and CTF refinement were carried out in RELION 3.0.4 (Zivanov et al., 2018).

In case of activator bound to TRPC4 bound structure, the refinement did not improve above 4.1 Å, as the dataset collected with AFIS suffered from stronger beam tilt which was estimated and corrected in RELION before 3D classification. For both the ligands, a 3D classification was performed with C4 symmetry to classify the subpopulation. The classes having high resolution features bound with ligands were selected and further polished and CTF-refined in RELION.

For TRPC4 bound with CaM, 3D classification using Sort3d in SPHIRE was performed to identify subpopulations with different complex stoichiometries. To further improve the resolution of the CaM region, we used symmetry expansion by quadrupling the 227,693 particles to mimic the C4 symmetry of the tetramer. Thus, the resulting 910,772 particles were used for Sort3d with a focused mask comprising the four CaM regions without imposing symmetry. Ten different classes obtained with Sort3d showed different stoichiometries (TRPC4 monomer:CaM) as shown in Figure S10. Four classes showing well resolved helices for CaM were selected and oriented in the same direction in order to boost the density at single CaM site (Figure S10). This rotation was achieved by applying a rotation of (± 90°, 180°, 270°) to the projection parameters of the classes using a customized script. After rotation, duplicates were removed, reducing the number of particles to 160,829. These particles were further polished and CTF-refined in RELION. The polished particles were finally refined in MERIDIEN (SPHIRE) with C1 symmetry using a mask encompassing TRPC4 with a single CaM.

### Local resolution estimation and filtering

The final half-maps were combined using a tight mask with the application of B-factors automatically determined by the PostRefiner tool in SPHIRE and filtered to the estimated resolution. The final estimated resolution by the ‘gold standard’ FSC = 0.143 criterion between the two masked half-maps is given in Table 2. The local resolution was calculated using sp_locres in SPHIRE. In case of TRPC4-CaM, the final densities were filtered according to local resolution using the local de-noising filter LAFTER (Ramlaul et al., 2019) to recover features with more signal than noise (based on half-maps).

**Table 2.**
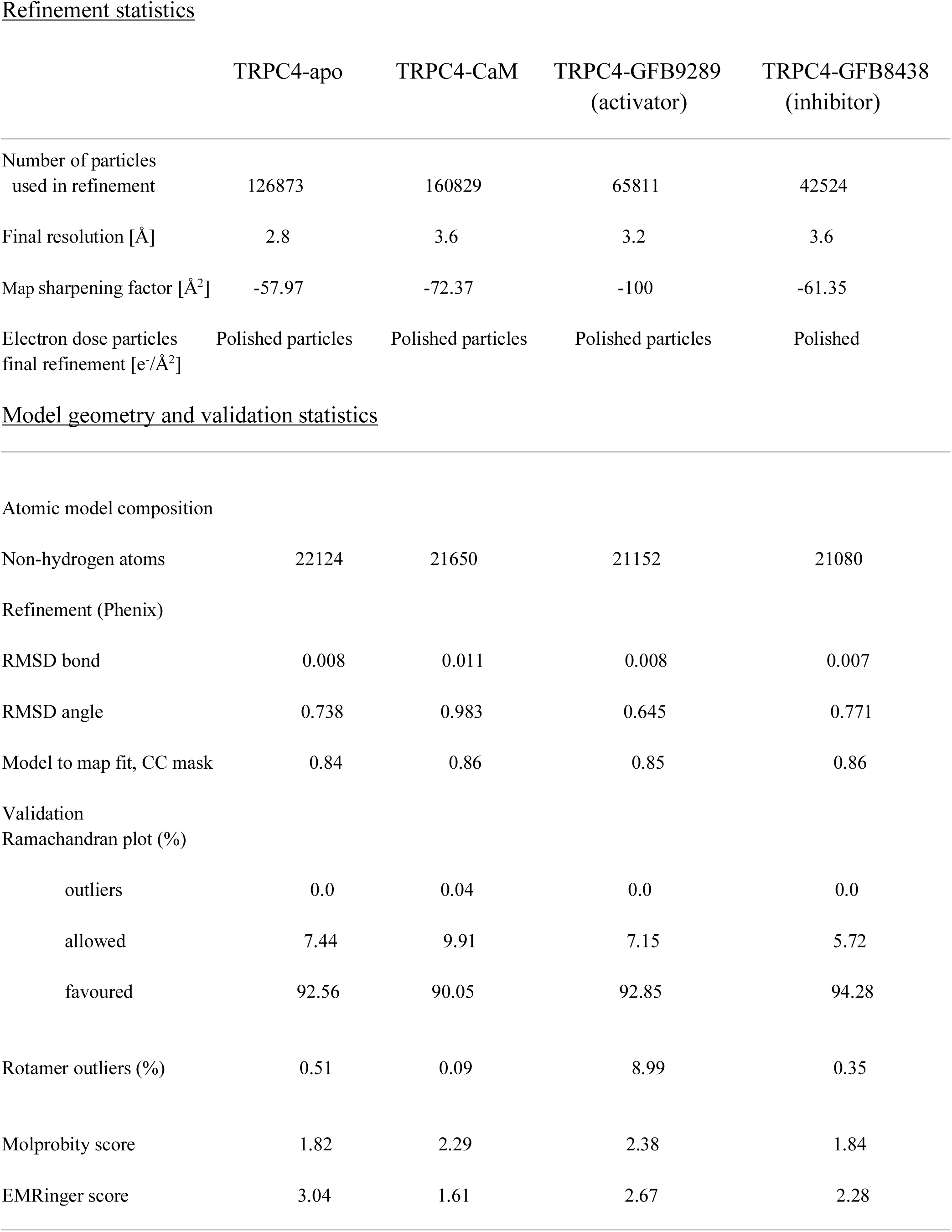
Refinement and model validation statistics.

|  | TRPC4-apo | TRPC4-CaM | TRPC4-GFB9289<br>(activator) | TRPC4-GFB8438<br>(inhibitor) |
| --- | --- | --- | --- | --- |
| Number of particles<br>used in refinement | 126873 | 160829 | 65811 | 42524 |
| Final resolution [Å] | 2.8 | 3.6 | 3.2 | 3.6 |
| Map sharpening factor [Å<sup>2</sup>] | -57.97 | -72.37 | -100 | -61.35 |
| Electron dose particles<br>final refinement [e/Å<sup>2</sup>] | Polished particles | Polished particles | Polished particles | Polished |
| Atomic model composition |  |  |  |  |
| Non-hydrogen atoms | 22124 | 21650 | 21152 | 21080 |
| Refinement (Phenix) |  |  |  |  |
| RMSD bond | 0.008 | 0.011 | 0.008 | 0.007 |
| RMSD angle | 0.738 | 0.983 | 0.645 | 0.771 |
| Model to map fit, CC mask | 0.84 | 0.86 | 0.85 | 0.86 |
| Ramachandran plot (%) |  |  |  |  |
| outliers | 0.0 | 0.04 | 0.0 | 0.0 |
| allowed | 7.44 | 9.91 | 7.15 | 5.72 |
| favoured | 92.56 | 90.05 | 92.85 | 94.28 |
| Rotamer outliers (%) | 0.51 | 0.09 | 8.99 | 0.35 |
| Molprobity score | 1.82 | 2.29 | 2.38 | 1.84 |
| EMRinger score | 3.04 | 1.61 | 2.67 | 2.28 |

### Model building, refinement and validation

The previously reported model of TRPC4 (Vinayagam et al., 2018) was initially docked into the density and fitted into the map as rigid body using UCSF Chimera. The model was further adjusted to fit in the density using Coot (Emsley et al., 2010) with an iterative process of real space refinement in Phenix (Adams et al., 2010) and model adjustment in Coot until a convergence evaluated by model to map fit with valid geometrical parameters. The high resolution obtained with the activator and apo structure enabled accurate modelling of the structure especially in the region encompassing residues 727-731 that connects the rib helix to the C-terminal helix (Figure S11). The presence of connecting density at this region shows the swapping of helix in this region. In our previous model, the density for the corresponding area was less resolved and the C-terminal helix was modelled without the domain swapping of the C-terminal helix. For both the activator and inhibitor molecules, cif files were generated using eLBOW tool in Phenix and used as geometrical restrains in Coot and Phenix during modelling and refinement respectively.

In the TRPC4-CaM complex, the C-terminal lobe of CaM bound with myosin light chain kinase (PDB ID: 2LV6) was used initially for rigid body fitting into the CaM density using Chimera. Then CaM was flexibly fitted into the density with the Cryo_fit tool in Phenix employing MD simulations. With the guide of this CaM model, the CaM was further adjusted manually to fit inside the density using Coot. Several rounds of iterative model building and refinement were performed using Coot and Phenix respectively until a good fit with a valid geometry was obtained (Table 2).

The densities corresponding to annular lipids were modelled as phosphatidic acid lipid (PDB ligand ID LPP) in the structures of activator-bound TRPC4 and apo TRPC4. In case of CaM-bound TRPC4 and inhibitor-bound TRPC4, a shorter lipid tail (PDB ligand ID 44E) was modelled due to limited resolution.

Finally, validation statistics computed by Phenix using MolProbity (Chen et al., 2010) were used to validate the overall geometry of the model, the model-to-map correlation value to assess the fitness of the model to its density, and an EMRinger score (Barad et al., 2015) to validate side chain geometry.

Figures were prepared in Chimera (Pettersen et al., 2004). Multiple sequence alignment was done using Clustal Omega (Sievers et al., 2011). Figures of the sequence alignment were made in Jalview (Waterhouse et al., 2009). The radius of the TRPC4 pore was determined using HOLE (Smart et al., 1996).

### Synthesis of GFB-9289

#### 4-chloro-5-(4-cyclohexyl-3-oxopiperazin-1-yl)-2,3-dihydropyridazin-3-one (GFB-9289)

To a solution of 1-cyclohexylpiperazin-2-one (150 mg, 0.8 mmol, 1 equivalent) in DMF (5 mL) was added 4,5-dichloro-2,3-dihydropyridazin-3-one (410 mg, 2.5 mmol, 3.0 equivalent) and DIEA (442 mg, 3.4 mmol, 4.0 equivalent) at ambient temperature under air atmosphere. The resulting mixture was stirred for 5 h at 100 °C. Then the reaction mixture was cooled and purified by reverse phase flash with the following conditions (Column: C18 OBD Column, 5um, 19×330mm; Mobile Phase A: Water (5 mmol/L NH_4_HCO_3_), Mobile Phase B: ACN; Flow rate: 45 mL/min; Gradient: 30% B to 60% B in 40 min; 254 nm; Rt: 15min) to afford crude product (80 mg), which was further purified by Chiral-Prep-HPLC with the following conditions: Column: CHIRALPAK IG-3, Column size :0.46×5cm;3um; Mobile phase: Hex(0.1%DEA):EtOH=80:20; Pressure: MPA; Flow: 1.0ml/min; Instrument: LC-08; Detector: 254nm; Temperature: 25 °C. 4-chloro-5-(4-cyclohexyl-3-oxopiperazin-1-yl)-2,3-dihydropyridazin-3-one (26.5 mg, 10.4%) was obtained at 1.436 min as a white solid (26.5 mg). ^1^H NMR (400 MHz, DMSO-*d*_6_) chemical shifts *δ* 12.91 (s, 1H), 7.86 (s, 1H), 4.23 (t, *J* = 12.1 Hz, 1H), 4.09 (s, 2H), 3.68 (t, *J* = 5.2 Hz, 2H), 3.38 (t, *J* = 5.3 Hz, 2H), 1.77 (d, *J* = 12.8 Hz, 2H), 1.61 (d, *J* = 15.6 Hz, 2H), 1.58 ‥C 1.40 (m, 3H), 1.31 (q, *J* = 13.1 Hz, 2H), 1.11 (t, *J* = 13.1 Hz, 1H). LRMS (ESI) *m/z*: [M+H]^+^ calculated for C_14_H_20_ClN_4_O_2_ 311.13; found 311.15. Purity 96%.

### Cellular Ca^2+^ uptake assay

Two 96-well plates of HEK293T cells had wells individually seeded at a density of 0.01 x 10^6^ cells in 200 µL DMEM/F12 + 10% FBS media per well, and were then grown in a 37 °C incubator with a 5% CO_2_ atmosphere. After 24 h, each well of one plate was individually transfected with the TRPC4_DR_ plasmid using Lipofectamine (Thermo Fisher Scientific). The other plate was left untransfected and used as a control. After 24 h, the transfection efficiency was confirmed by monitoring fluorescence of the EGFP fused to TRPC4_DR_ using an EVOS FL microscope (Thermo Fisher Scientific).

A 1 mM working solution of rhod-2 AM ester was prepared by mixing 25 µL DMSO + 25 µL of 20% Pluronic F-127 per 50 µg rhod-2 AM ester. 50 µL of rhod-2 AM ester loading buffer (10 µM rhod-2 AM ester in working solution, 10 mM HEPES pH 7.4, 130 mM NaCl, 5 mM KCl, 8 mM glucose, 1.2 mM MgCl_2_, 1.5 mM CaCl_2_, 0.05% Pluronic F-127) was then added to each well and incubated for 1.5 h at 37 °C. The media-loading buffer mix was thoroughly removed and 50 µL recording buffer (10 mM HEPES pH 7.4, 130 mM NaCl, 5 mM KCl, 8 mM glucose, 1.2 mM MgCl_2_, 1.5 mM CaCl_2_, 0.05% Pluronic F-127) was added per well. Rhod-2 fluorescence of the cells was recorded using a Spark multimode microplate reader (Tecan) prior to each experiment. Rhod-2 uptake efficiency was assumed to be similar across all wells, and these values were used as a proxy for the number of cells per well, a parameter that was utilized later during data processing.

The following conditions were tested in order to determine the effect of agonists and antagonists on Ca^2+^ influx into TRPC4_DR_- and non-transfected cells: nothing added (additional negative control), + 0.28% DMSO (negative control), + 112 nM Englerin A (positive control), + 112 nM Englerin A / 16800 nM GFB-8438, + 10630 nM GFB-9289, + 10630 nM GFB-9289 / 16800 nM GFB-8438. 16 wells containing biological replicates were used per condition tested per plate. Except for the condition where no extra components were added, compounds at 2x final concentration in a total of 50 µL recording buffer were added per well. In conditions where the antagonist GFB-8438 was used, the cells were pre-incubated for 5 min with 25 µL GFB-8438 at 4x final concentration before adding 25 µL of Englerin A or GFB-9289 at 4x final concentration and initiating recording. Recording of rhod-2 fluorescence increase (indicative of Ca^2+^ influx into cells) immediately after addition of each condition’s final component was performed on a Spark multimode microplate reader (Tecan) at an excitation wavelength of 550 nm and an emission wavelength of 581 nm. Recording occurred for 1 minute over 10 cycles with 6 seconds per cycle.

During data analysis, the rhod-2 fluorescence signal of cycle 1 was subtracted for every well in order to obtain a baseline value. Normalization coefficients for cell numbers were calculated based on the rhod-2 fluorescence measured prior to compound addition and applied to the analyzed data, allowing the results of the assay to be compared between the biological replicates. Biological replicates that were subject to handling and technical errors during the measurement were considered outliers and excluded. The resulting data was plotted, which utilized the following number of biological replicates (TRPC4-transfected / non-transfected cells): 15/16 for nothing added, 11/12 for + DMSO, 10/14 for + Englerin A, 12/15 for + Englerin A / GFB-8438, 10/12 for + GFB-9289, 13/12 for + GFB-9289 / GFB-8438.

## Acknowledgements

We are thankful to Nina Ludwigs, Marion Hülseweh and Nathalie Bleimling for technical assistance. We also thank Ingrid Vetter and Sabrina Pospich for helpful discussion with model building and analysis and Amrita Rai for the assay development. We are grateful to Matthew Daniels for assistance with compound comparison and selection. This work was supported by funds from the Max Planck Society (to S.R.).

## Data availability

The atomic coordinates and cryo-EM maps for TRPC4_DR_ in complex with XXX, XXX are available at the Protein Data Bank (PDB)/Electron Microscopy Data Bank (EMDB) databases. The accession numbers are XXXX/EMD-XXXX. The data sets generated in the current study are available from the corresponding author on reasonable request.

## Author Contributions

S.R. and G.M. designed the project. D.V. expressed and purified TRPC4. O.H. recorded the EM images. D.V. prepared cryo-EM specimens, D.V. and F.M. and M.S. processed cryo-EM data, and D.V. built the atomic models. O.S. performed the Ca^2+^ uptake experiments. M.Y. and M.W.L. designed and executed compound synthesis. G.M. implemented QC assay data. D.V., O.S. and D.Q. prepared figures and movies. D.V., D.Q. and S.R. wrote the manuscript. All authors reviewed the results and edited the manuscript.

---

**Figure S1.**
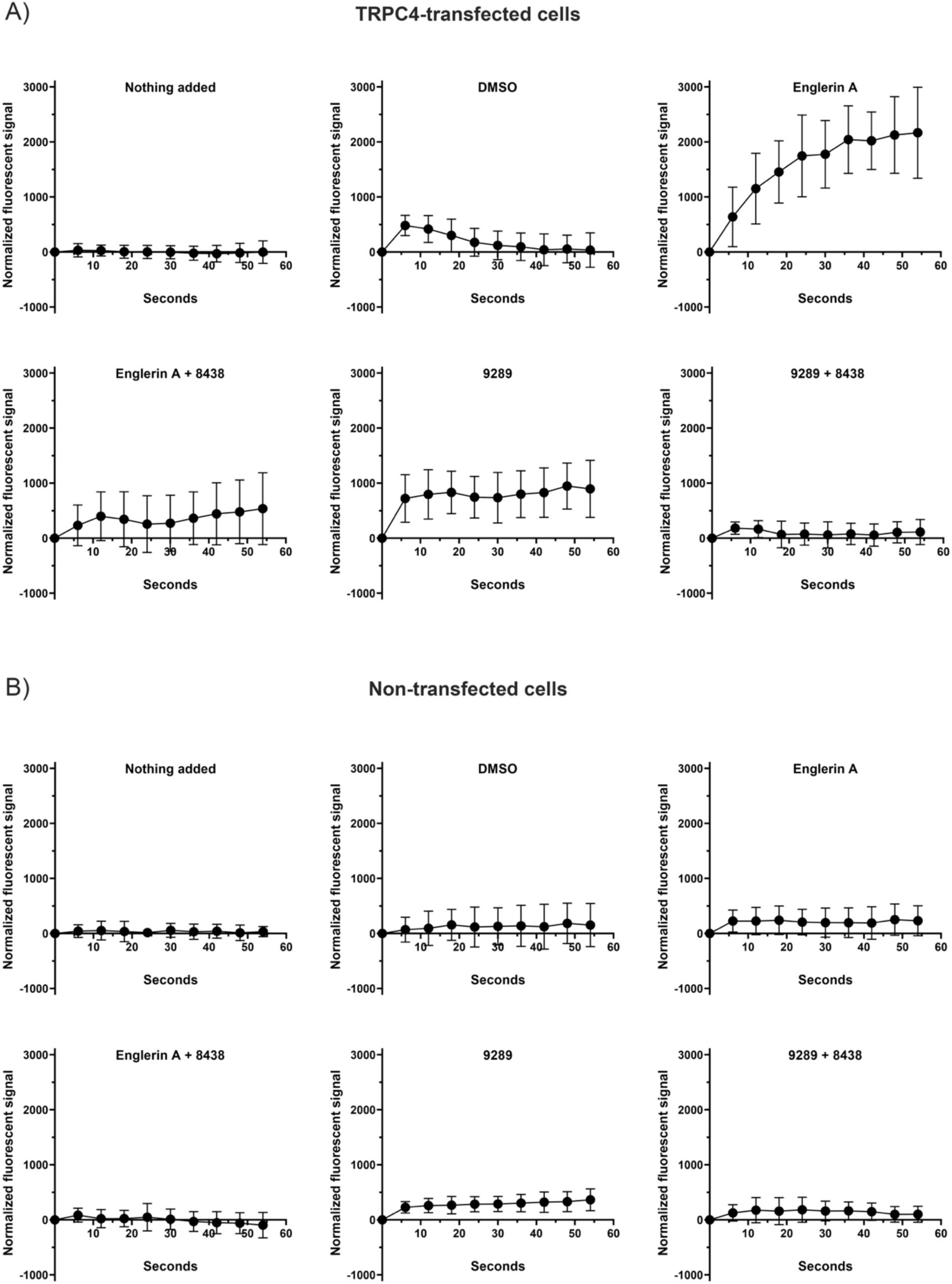
Effect of TRPC4 activators and inhibitors on Ca^2+^ uptake in cells. (A) The effect of TRPC4 activators and inhibitors in a cellular context was assessed by transfecting HEK293T cells with TRPC4, loading them with the Ca^2+^-sensing dye rhod-2, and assessing the fluorescence at 581 nm of rhod-2 after incubation with either controls or TRPC4 activity-modulating compounds. Cells treated with the potent TRPC4 activator Englerin A show a significant increase in rhod-2 fluorescence indicative of strong Ca^2+^ influx. Incubation with GFB-9289 also results in activation of the channel, albeit at a more modest level than Englerin A. Prior incubation with GFB-8438 significantly decreases Ca^2+^ influx into cells, when activated by Englerin A or GFB-9289, highlighting is inhibitory function. (B) The same experiment shown in (A) was also performed on non-transfected cells, demonstrating that the effect of the tested activator and inhibitor on Ca^2+^ influx was mainly due to modulation of transfected TRPC4 activity.

**Figure S2.**
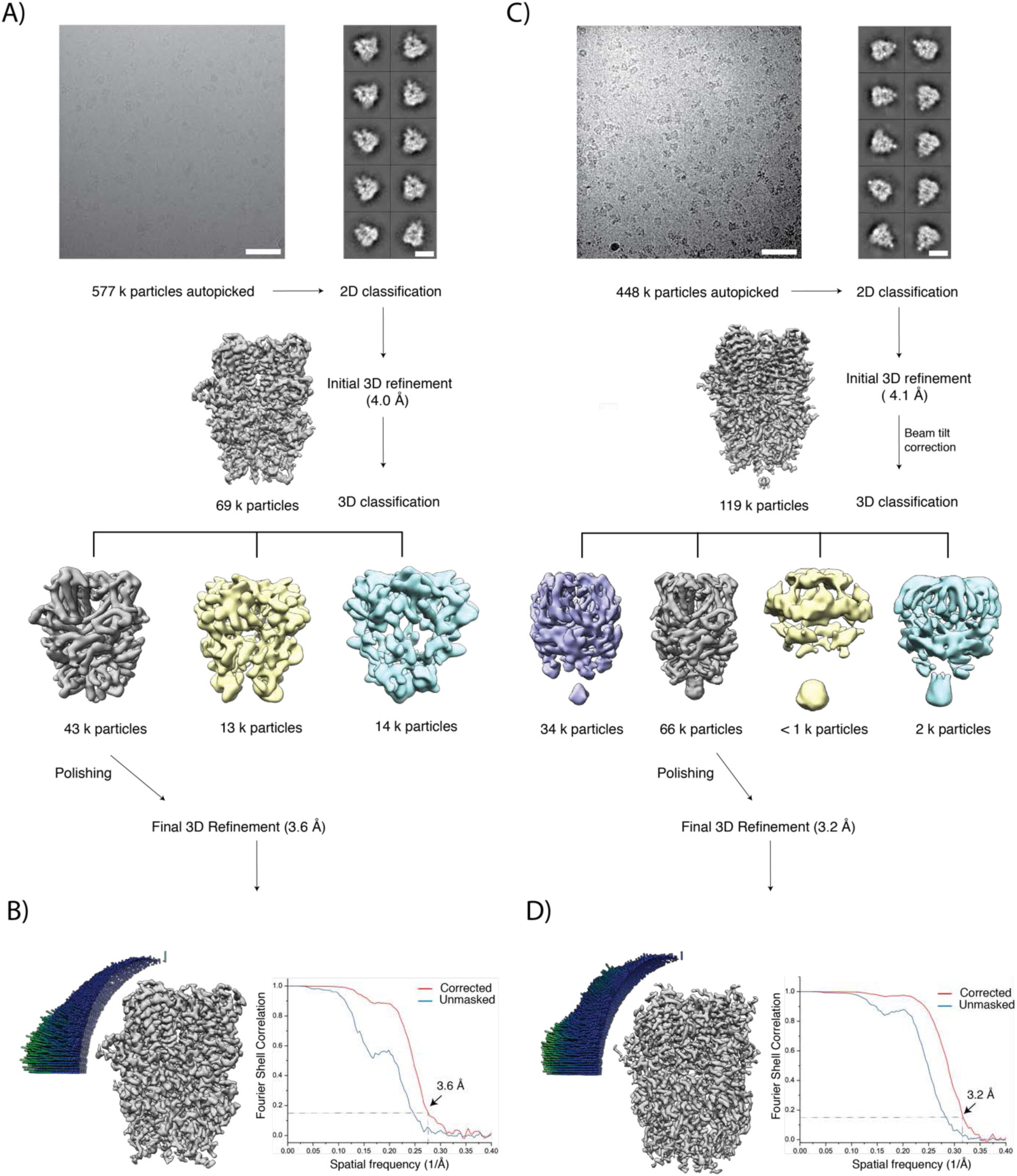
Cryo-EM image processing workflow for TRPC4 in complex with the inhibitor or activator. **(A)** The top panel shows a representative digital micrograph area and selected 2-D class averages of inhibitor bound TRPC4. Scale bars, 50 nm and 10 nm, respectively. The initial refinement density and subsequent densities obtained after 3D classification are shown in the middle and bottom panel, respectively. **(B)** Angular distribution of particles used in the final refinement and Fourier shell correlation curves (FSC) between the two independently refined maps. The dotted lines indicate the 0.143 FSC criterion used for average resolution estimation. **(C, D)** Same as in (A) and (B), respectively, but for the activator-TRPC4 complex.

**Figure S3.**
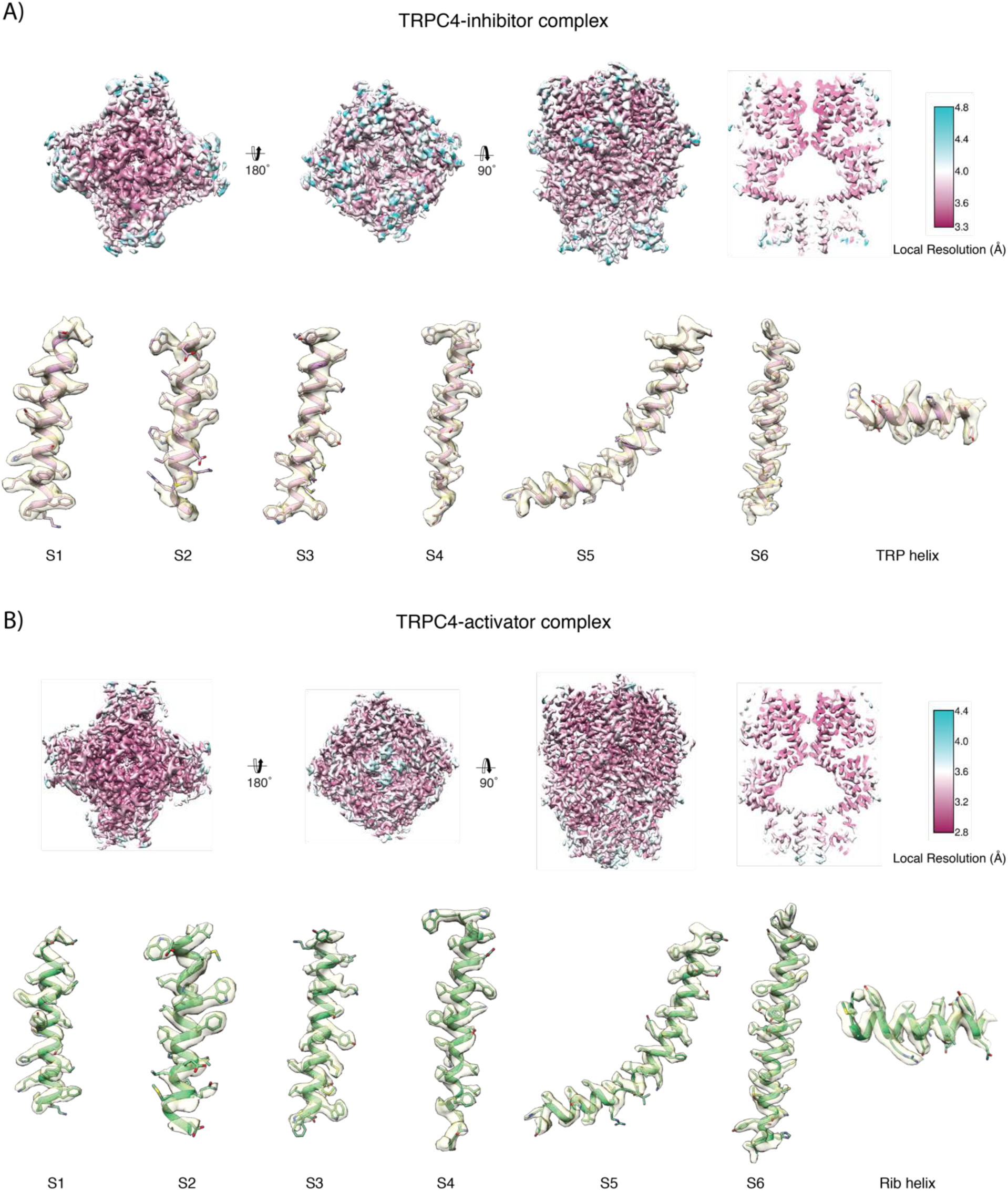
Local resolution of the TRPC4-ligand complex maps. **(A, B)** Maps of inhibitor-(GFB-9289) and activator (GFB-8348)-bound TRPC4, respectively, coloured according to the local resolution. Representative regions of the density with the fitted atomic model are shown below the local resolution maps.

**Figure S4.**
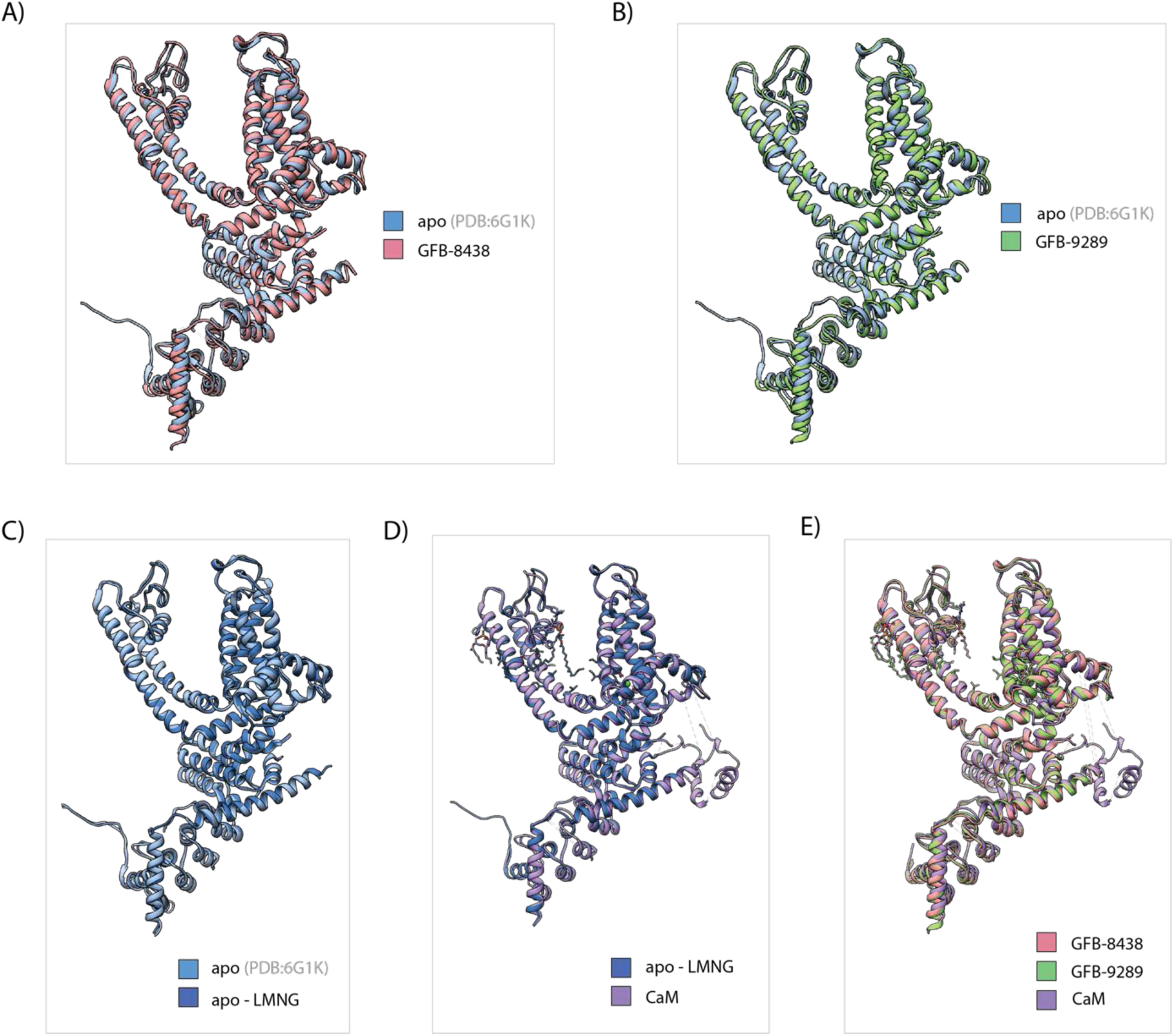
Comparison between different TRPC4 structures. (**A**) Structural alignment of a protomer of the inhibitor bound TRPC4 with that of TRPC4 in its apo state. The protomer of the TRPC4 apo structure is shown in cartoon representation and colored in light blue while the protomer of the inhibitor bound TRPC4 structure is colored in light red. (**B**) Same as in **(A)** for the activator bound TRPC4. The protomer of activator bound TRPC4 structure is shown in light green. (**C**) Alignment of the structures of TRPC4 solubilized in amphipols or in the detergent LMNG. The protomer of apo-LMNG structure is shown in dark blue. (**D**) Alignment of the TRPC4-CaM complex structure with the TRPC4 apo-LMNG structure. The TRPC4-CaM complex is also solubilized in LMNG. The protomer of the TRPC4-CaM structure is shown in purple. (**E**) Alignment of the TRPC4-activator, TRPC4-inhibitor and TRPC4-CaM structures. Note: The C-terminal helix in 6GIK pdb has been corrected for domain swapping (See Method section and Figure S12 for details)

**Figure S5.**
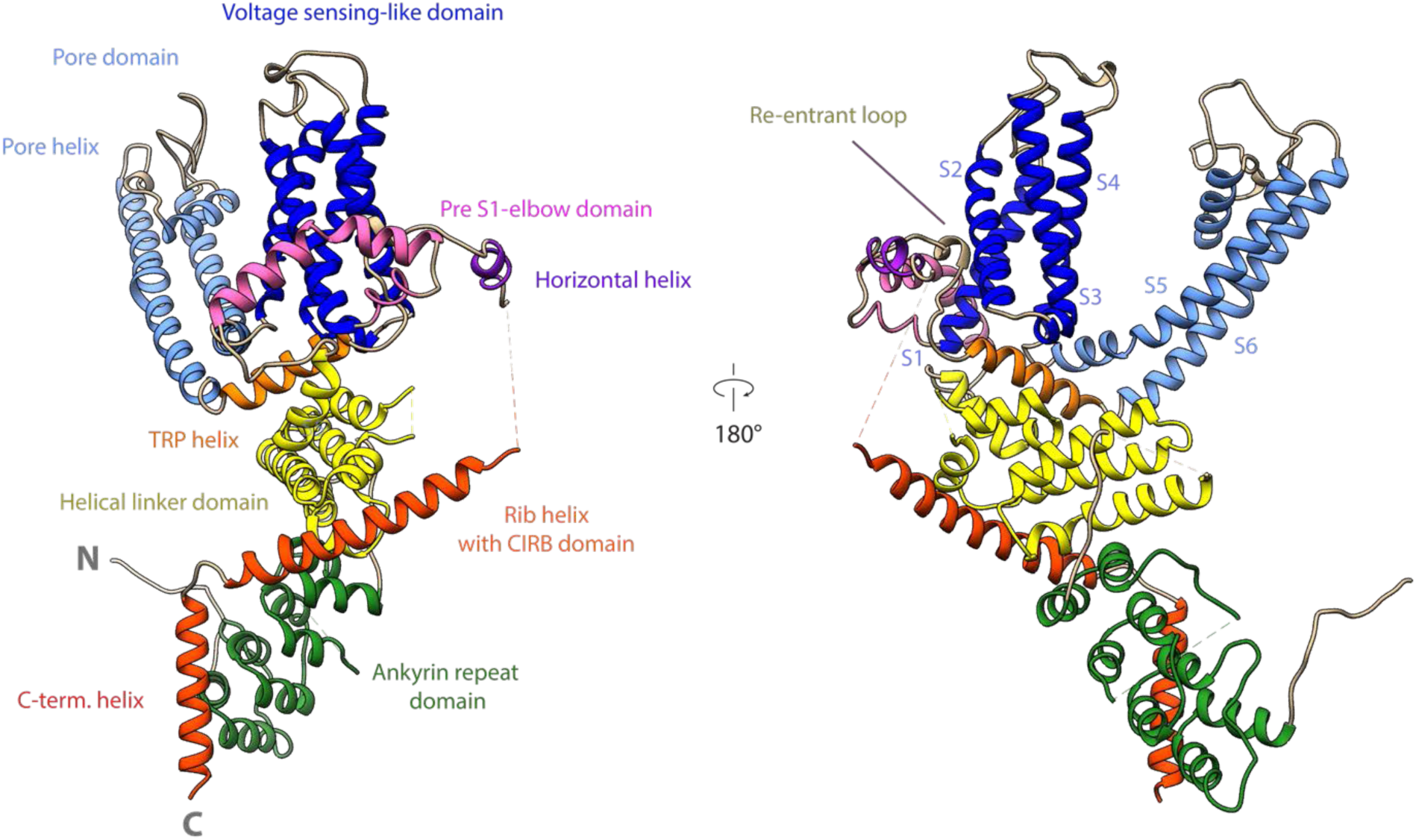
Domain architecture of zebrafish TRPC4 channel. Cartoon representation of a TRPC4 protomer. Each domain is shown in a different color and labeled accordingly.

**Figure S6.**
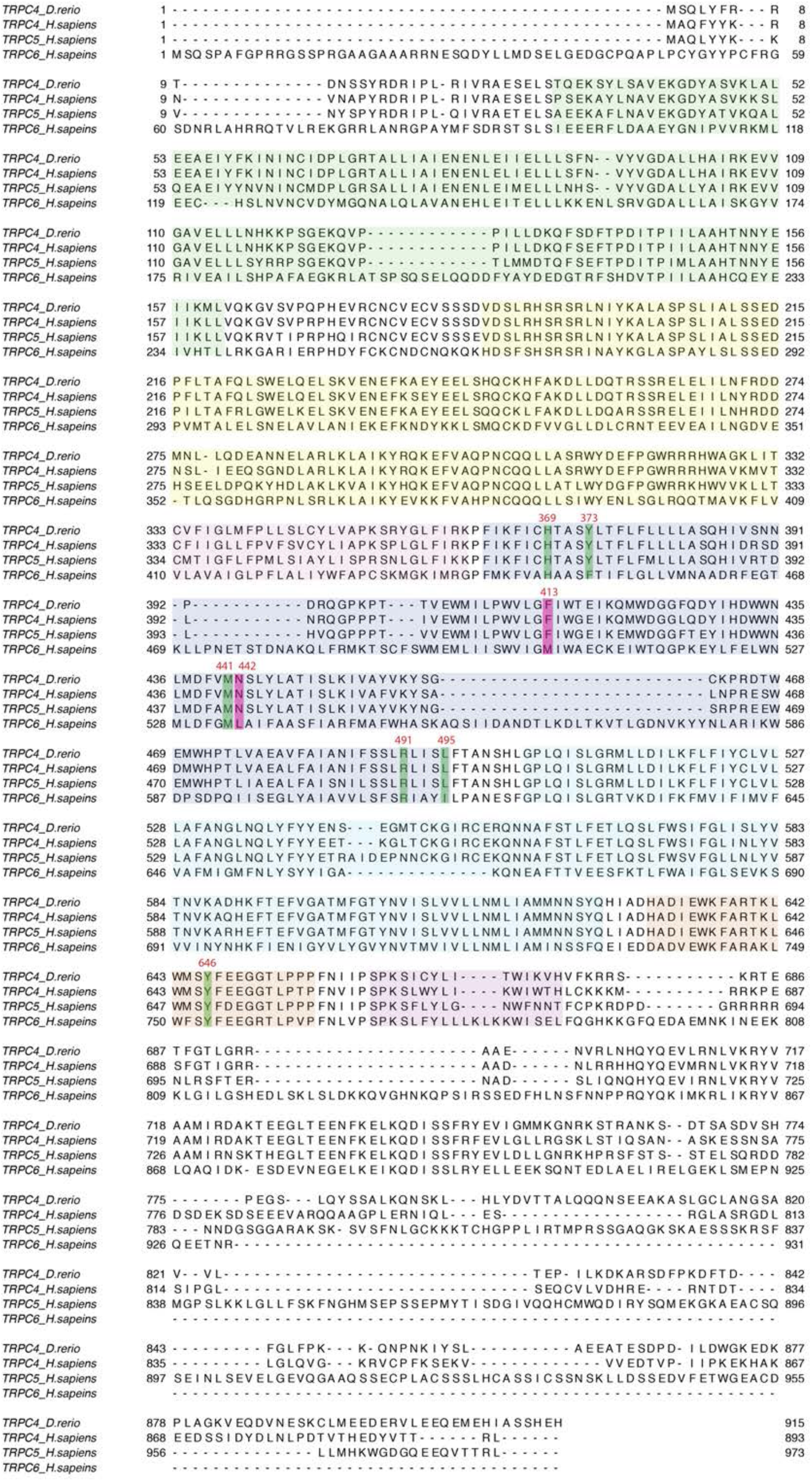
Sequence alignment of zebrafish TRPC4, human TRPC4, TRPC5, and TRPC6. The highlighted and marked residues denote the conserved residues in TRPC4 and TRPC5 interacting with the inhibitor GFB 8438. The residues highlighted in red colour shows the critical difference between TRPC4/5 and TRPC6 for the inhibitor binding site.

**Figure S7.**
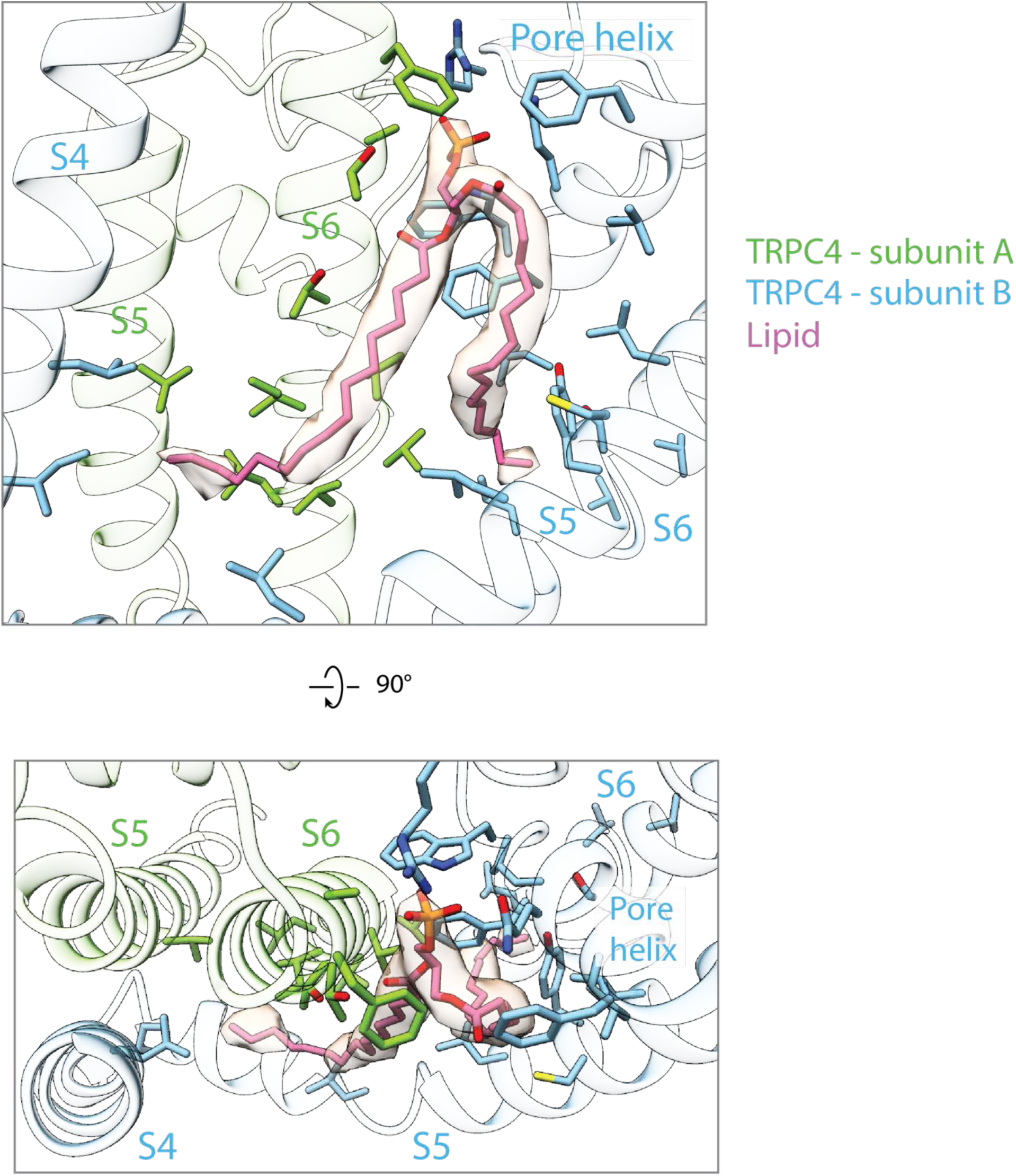
Different views of the lipid binding pocket at the interface between two subunits. Phosphatidic acid (pink) that binds at the interface between two subunits near the pore region is shown in stick representation along with the corresponding density. The interacting residues from the S4, S5 and S6 helices are also shown in stick representation. The protein residues from different protomers are colored differently. The helices are shown in cartoon representation with high transparency.

**Figure S8.**
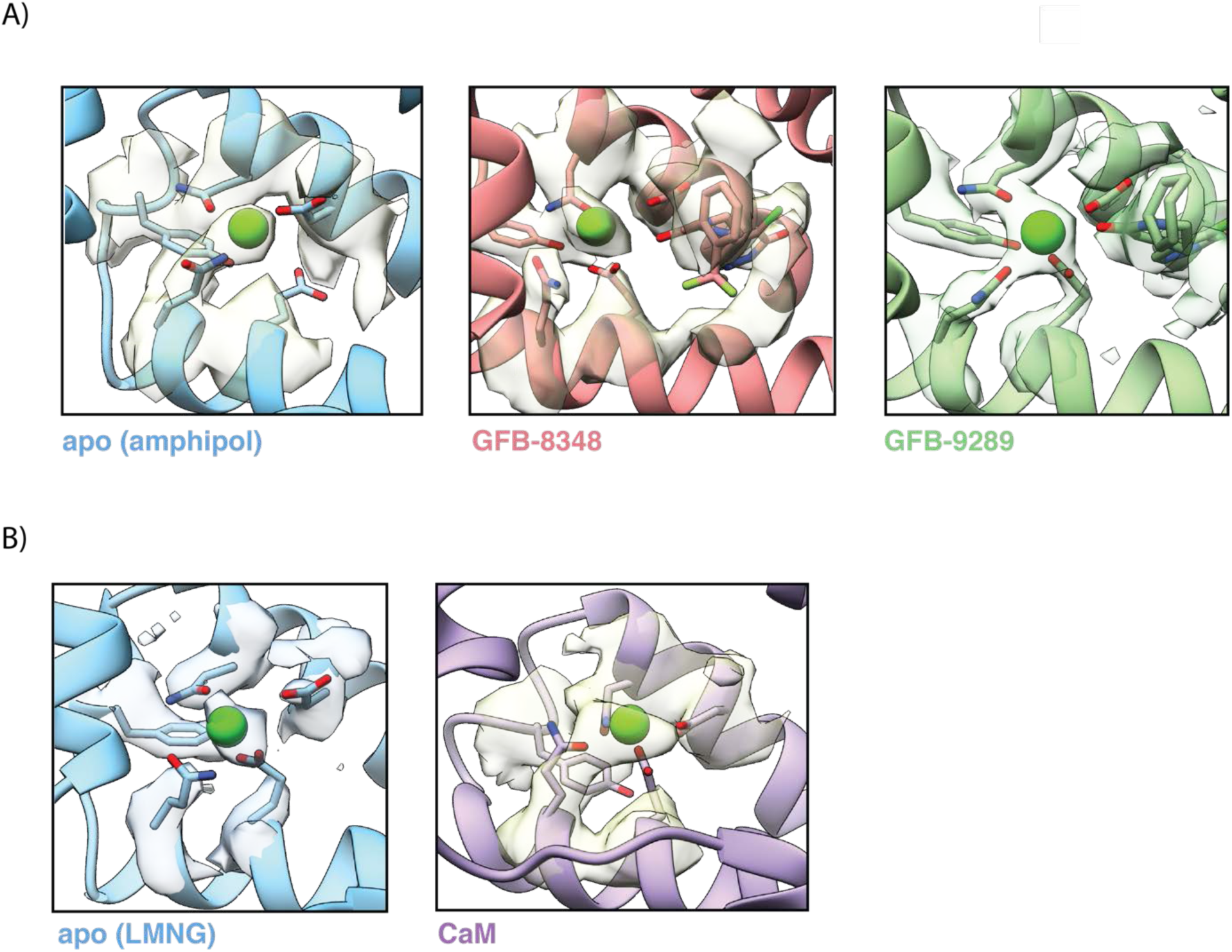
Ca^2+^-binding site in the VSL domain of apo and ligand bound TRPC4. **(A)** Close-up view of the Ca^2+^-binding site in apo and ligand bound TRPC4 determined in amphiphols. The coordinating residues in the Ca^2+^ ion binding site and the modelled Ca^2+^ ion are shown in stick and sphere representation along with their densities. The oxygen atom of the ligand molecules is situated close to the bound Ca^2+^-ion. **(B)** Close-up view of the Ca^2+^-binding site in apo and TRPC4-CaM complex determined in presence of Lauryl Maltose Neopentyl Glycol (LMNG). The external addition of Ca^2+^ ion during the TRPC4-CaM complex preparation reflects its strong density.The different structural forms shown in (A) and (B)are color coded and named accordingly.

**Figure S9.**
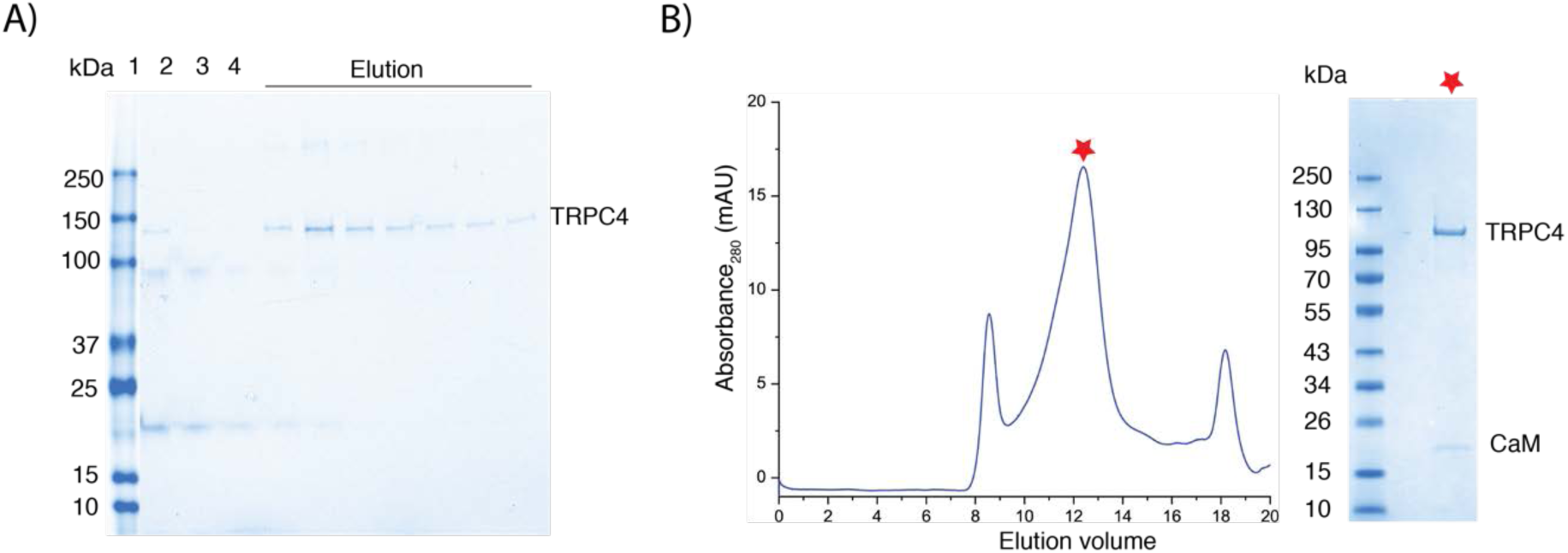
Analysis of CaM binding to TRPC4 by biochemical methods. **(A)** SDS gel electrophoresis analysis of the TRPC4 pull down experiment performed with a CaM Sepharose column. Lane 1 - protein size marker, lane 2 - TRPC4 input, lane 3 - flow through, lane 4 - wash, remaining lanes - elution fractions. **(B)** Gel filtration analysis of the TRPC4-CaM complex. The peak fraction containing the TRPC4-CaM complex are indicated by a star were further analysed by SDS gel electrophoresis (right panel).

**Figure S10.**
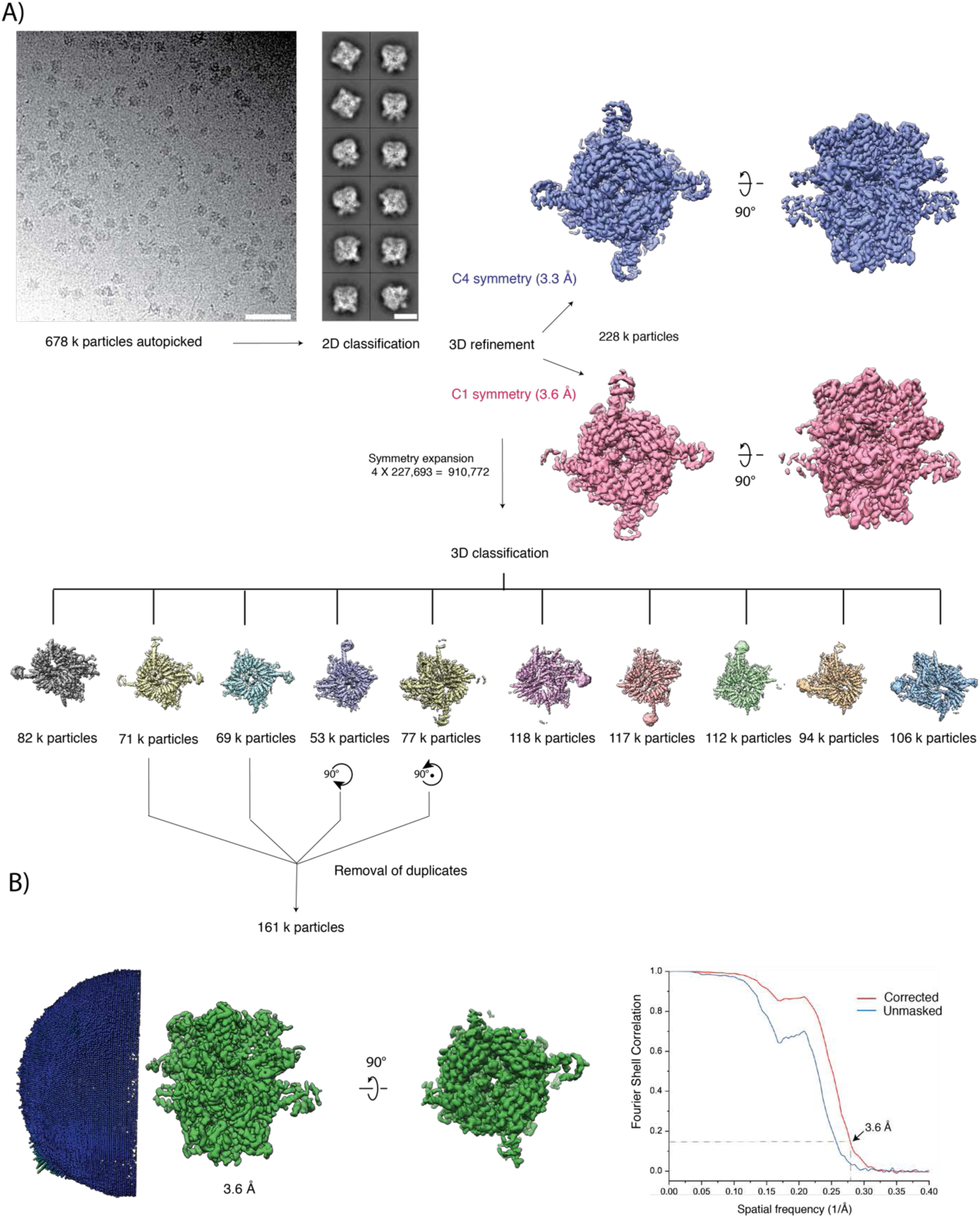
Cryo-EM image processing of the TRPC4-CaM complex. **(A)** The top left panels show a representative digital micrograph area and selected class averages of the TRPC4-CaM complex. Scale bars, 50 nm and 10 nm, respectively. The initial refinement densities obtained without symmetry (red) and with C4 symmetry (blue) are shown in the top right panel. The middle panel shows densities of different subclasses obtained with 3-D classification after symmetry expansion. Four subclasses were selected, rotated as indicated and used for the final refinement after removal of duplicates. **(B)** Angular distribution of particles used in the final refinement and Fourier shell correlation curves (FSC) between the two independently refined maps. The dotted lines indicate the 0.143 FSC criterion used for average resolution estimation.

**Figure S11.**
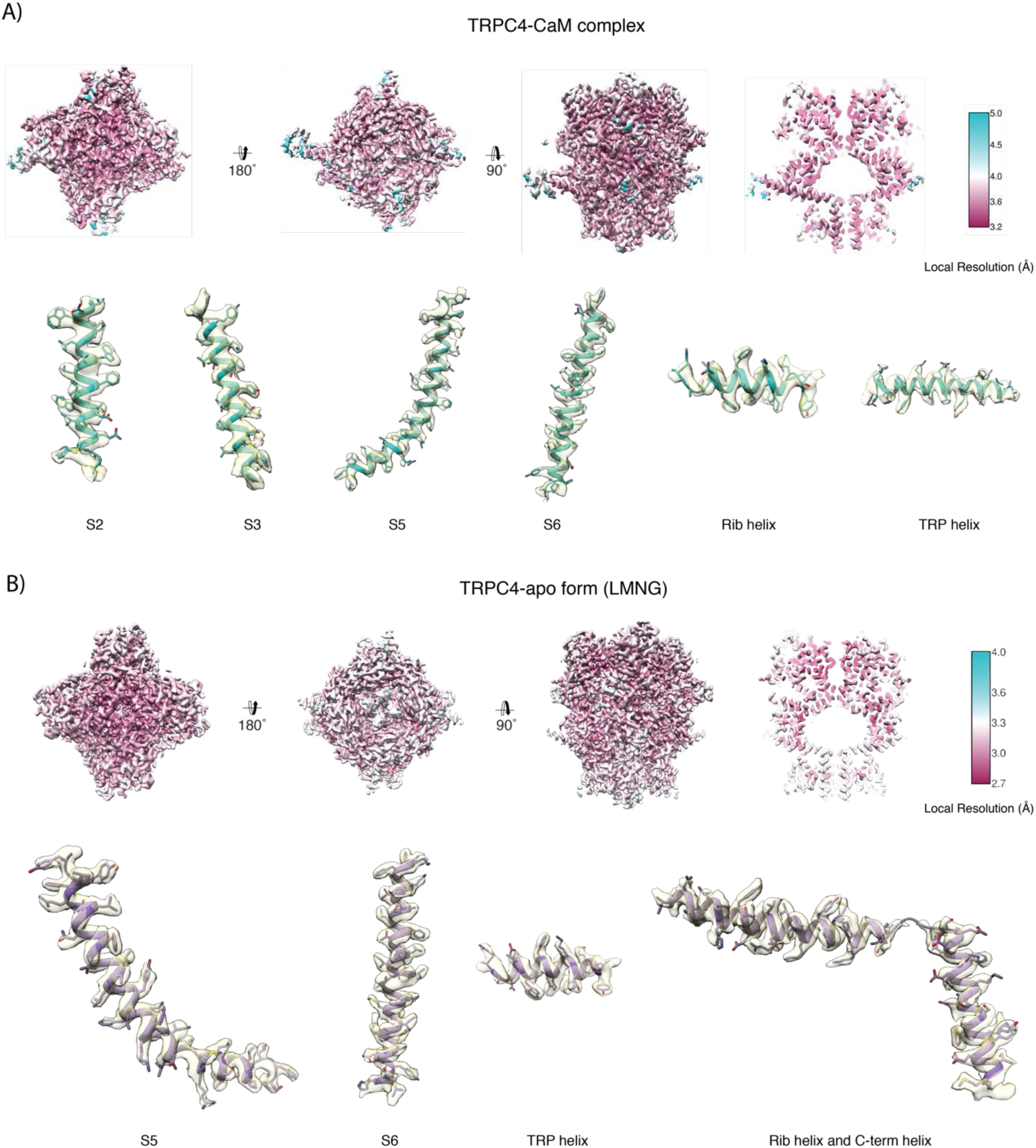
Local resolution maps of TRPC4-apo (LMNG) and TRPC4-CaM. **(A, B)** Maps of TRPC4-apo (LMNG) and TRPC4-CaM, respectively, coloured according to the local resolution. Representative regions of the density with the fitted atomic model are shown below the local resolution maps.

**Figure S12.**
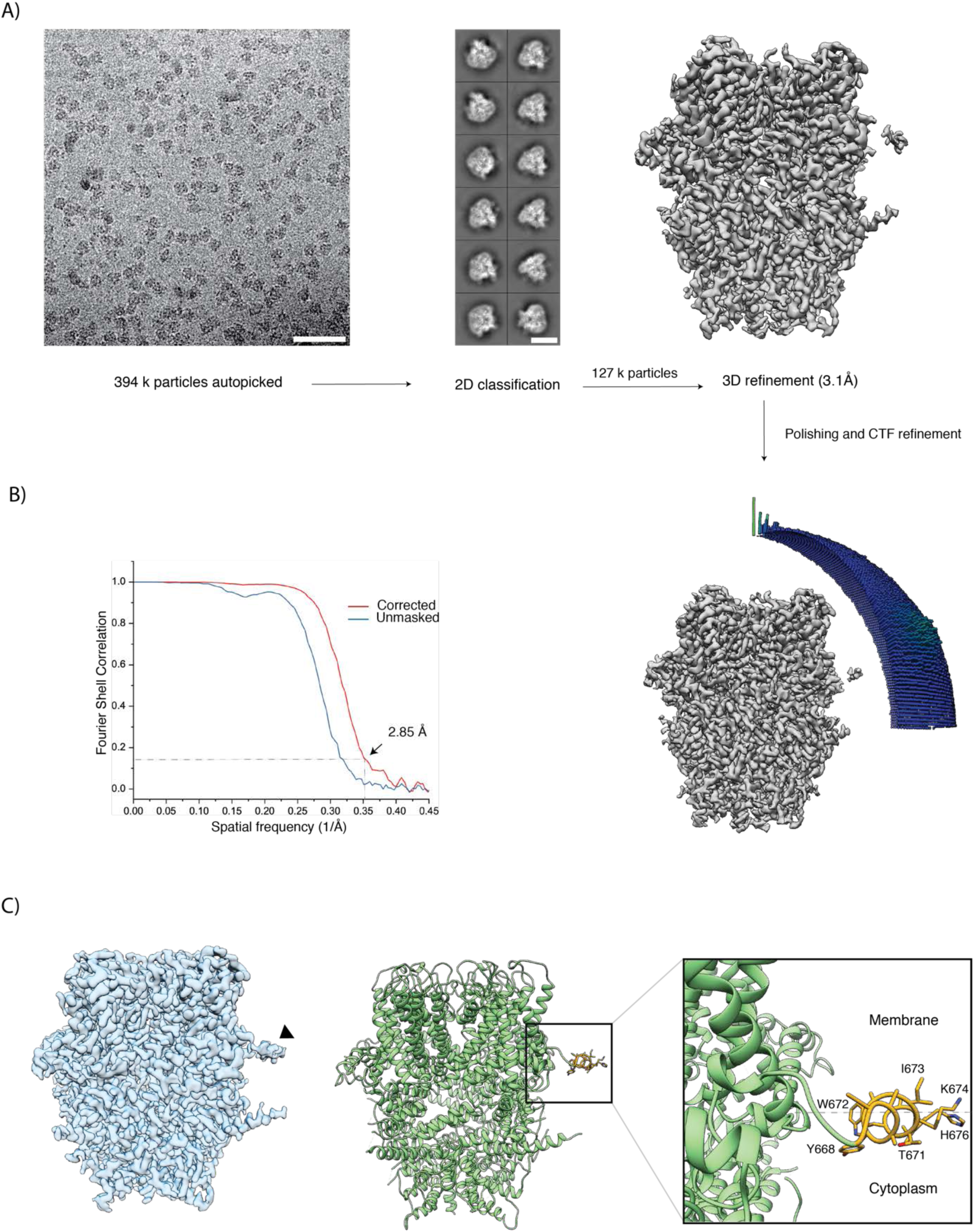
Cryo-EM image processing and structure determination of TRPC4 solubilized in LMNG. **(A)** The top left panels show a representative micrograph and class averages of TRPC4 solubilized in LMNG. Scale bars, 50 nm and 10 nm, respectively. The density resulting from the initial 3D refinement is shown in the top right panel. **(B)** Angular distribution of particles used in the final refinement and Fourier shell correlation curves (FSC) between the two independently refined maps. The dotted lines indicate the 0.143 FSC criterion used for average resolution estimation. **(C)** The final map, filtered using LAFTER, shows a clear density corresponding to a horizontal helix (indicated with an arrowhead) (left panel). The corresponding structure in cartoon representation with the residues of the horizontal helix shown in golden yellow (middle panel). Zoom-in view of the horizontal helix (right panel).

**Figure S13.**
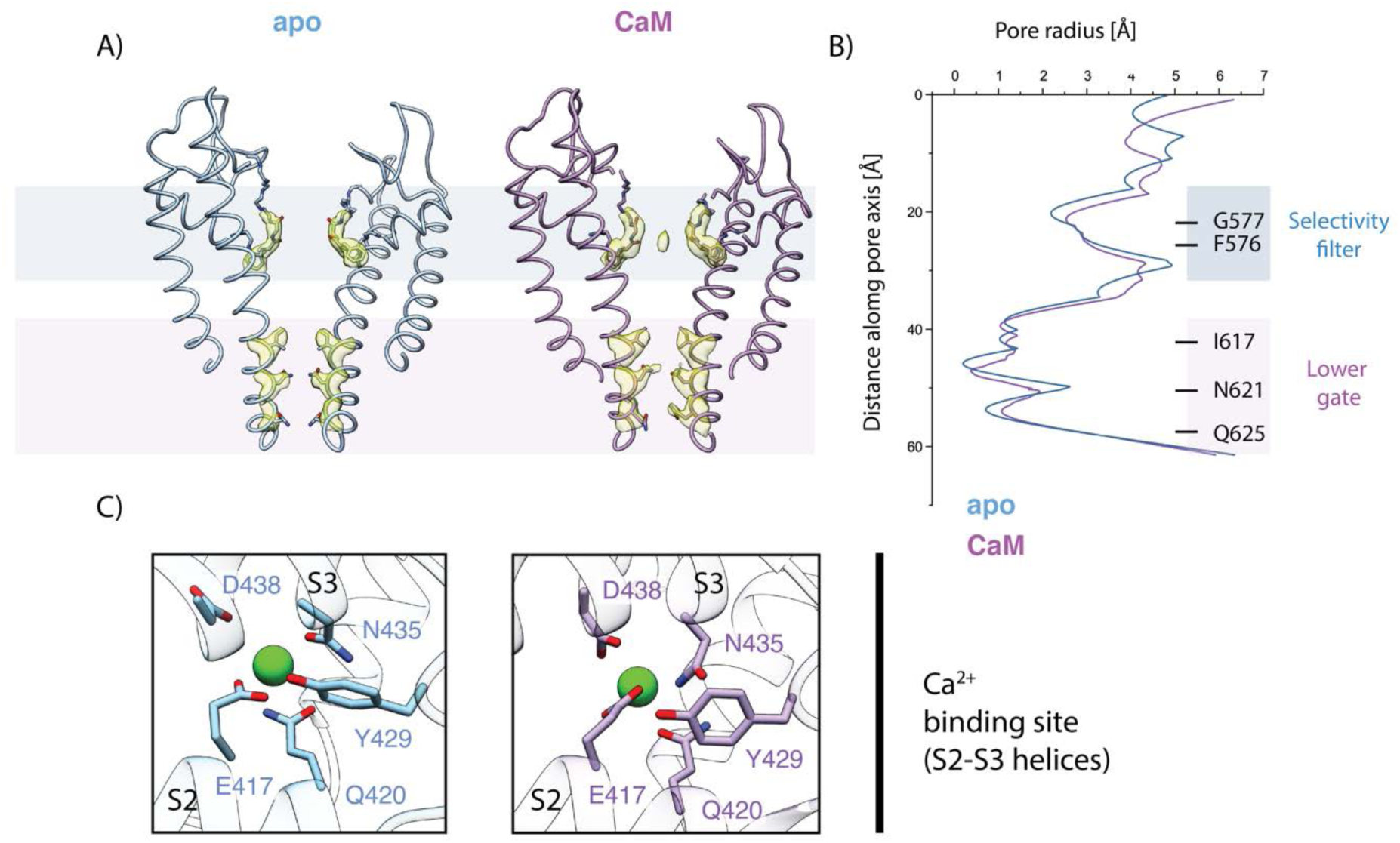
Comparison of the ion conduction pore and Ca^2+^-binding site. (**A**) Side view of the pore-forming region of TRPC4 in the apo (LMNG)- (blue), and CaM-bound (purple) structures. Only the two opposing subunits of the tetrameric channel are shown as ribbon representation for clarity. The density at comparable thresholds corresponding to the selectivity filter (light blue) and the lower gate (pink) is superimposed. Note, only in the TRPC4-CaM complex structure an additional density occupies the center of the selectivity filter. (**B**) The calculated pore radii corresponding to the structures in (A) are depicted. The color code is identical to (A). The positions of important residues, constituting the selectivity filter and the lower gate, are indicated on the right. (**C**) Close-up view of the Ca^2+^-binding site in the VSL domain of TRPC4-apo (LMNG) (left) and TRPC4-CaM (right). Ca^2+^ ion is shown as green sphere and interacting residues are highlighted. Color code of TRPC4 structures is as in (A).

